# CASTLE: Cell Adhesion with Supervised Training and Learning Environment

**DOI:** 10.1101/2020.06.02.130708

**Authors:** S G Gilbert, F Krautter, D Cooper, M Chimen, A J Iqbal, F Spill

## Abstract

Different types of microscopy are used to uncover signatures of cell adhesion and mechanics. Automating the identification and analysis often involve sacrificial routines of cell manipulation such as *in vitro* staining. Phase-contrast microscopy (PCM) is rarely used in automation due to the difficulties with poor quality images. However, it is the least intrusive method to provide insights into the dynamics of cells, where other types of microscopy are too destructive to monitor. In this study, we propose an efficient workflow to automate cell counting and morphology in PCM images. We introduce Cell Adhesion with Supervised Training and Learning Environment (CASTLE), available as a series of additional plugins to ImageJ. CASTLE combines effective techniques for phase-contrast image processing with statistical analysis and machine learning algorithms to interpret the results. The proposed workflow was validated by comparing the results to a manual count and manual segmentation of cells in images investigating different adherent cell types, including monocytes, neutrophils and platelets. In addition, the effect of different molecules on cell adhesion was characterised using CASTLE. For example, we demonstate that Galectin-9 leads to differences in adhesion of leukocytes. CASTLE also provides information using machine learning techniques, namely principal component analysis and *k*-means clustering, to distinguish morphology currently inaccessible with manual methods. All scripts and documentation is open-source and available at the corresponding GitLab project.

## 1. Introduction

The function of almost all cell types, including leukocytes, platelets, and cancer cells, critically involves a number of mechanical processes[1]; these include the regulation of cell-cell and cell-matrix adhesion, flow sensing and changes in morphology during cell migration[2]. A particularly important case, where adhesions need to dynamically change on short time scales, involve the adherence of blood cells. For example, leukocytes adhere to the endothelium and transmigrate out into inflamed tissues, and platelets activate and adhere to a substrate or to each other during wound closure. To investigate the effect of specific adhesion molecules on leukocyte behaviour, e.g. those appearing on the endothelium, biologists often employ flow based adhesion assays where cells adhere to coated substrates, or to endothelial monolayers. Subsequently, cells may migrate on the substrate or monolayer, or form aggregates. To identify adhesive states or morphological changes during migration, experimentalists frequently analyse a large number of cells under different conditions.

Humans are highly perceptive in identifying patterns in complex biological images. However, manual counting of the same object over hundreds of images is exhausting and time-consuming, increasing errors in the image analysis. Developments in computer power and algorithms therefore have great potential to allow more efficient routes to automatically process and analyse masses of data with limited human intervention if a suitable methodology can be put into place. Moreover, such automation can help to extract new quantitative features from the images.

High-throughput microscopy is the acquisition and processing of large volumes of data by automating sample preparation and data analysis techniques [3]. The methodology for these types of study are referred to as workflows. Workflows utilise the best combination of image analysis techniques, specific to a researcher’s problem, and follow the steps to ensure the method is repeated exactly for all samples. The primary objective of the methods chosen is in their accuracy for which features of an image are converted into meaningful quantitative data [4]. Once validated as accurate, designing a workflow with flexibility for similar research enables a larger impact. To be considered more widely usable by the community of bio-imaging researchers they need to excel in certain attributes. Carpenter et al. [5] list the criteria that such research software should meet. The criteria can be summarised into five key attributes: User-friendly, Modular, Developer-friendly, Validated and Interoperable. All of these must be considered for a successfully designed workflow.

In our study, we are considering the images acquired by PCM. Since most thin biological samples are optically transparent in visible light, amplitude information does not provide good contrast for imaging. However, even these transparent samples provide a significant optical phase delay. PCM utilises both types of information in the transmitted light, whereas conventional bright-field imaging only measures the amplitude. Full details of quantitative phase imaging techniques can be found elsewhere [6].

Once the images are uploaded to a digital form, the automated workflow processes the image into its constituent parts. There are many techniques in image processing for pixel classification [7, 8]. In addition, a methodology to interpret the vast amounts of statistics produced is necessary to prevent a backlog of valuable information going unused or slowing down the research. Again, techniques in machine learning can help automate the process with little human intervention to increase both speed and objectivity [9, 10].

In the present study, we test our workflow on different types of recently acquired data sets: we quantify adhesion and morphology of peripheral blood mononuclear cells (PBMCs) and polymorphonuclear leukocytes (PMNs) in dependence on galectins, and platelet adhesion depending on von Willebrand factor (vWF) with stimulation by a platelet agonist, Adenosine Diphosphate (ADP). Galectins are a family of ß-galactoside binding proteins which have a range of immunomodulatory functions [11]. More recently, Galectin-9 (Gal-9), a tandem repeat type galectin, has been proposed to play a role in modulating leukocyte trafficking [12]. However, its precise role in this process, in particular the way it regulates adhesion, remains elusive. Similarly, platelets need to adhere to substrates and to other platelets to perform their function. ADP is a known activator of platelets that switches on their GPIIb/IIIa receptors [13, 14], allowing them to bind to vWF that is present in the endothelium and subendothelial tissue [15]. Studying the response of adherent cells to stimuli such as ADP, or proteins such as vWF or Gal-9 that are involved in cell adhesions, is therefore critical to understanding cell adhesion.

This paper introduces a novel workflow to analyse images of adherent cells for images that are commonly obtained in experimental setups modelling blood cell recruitment within the vasculature towards sites of infections, chronic inflammation or wounds and coagulation. We demonstrate that our workflow is capable of identifying changes in cell adhesion as well as morphology depending on different experimental conditions; however, these adhesive and morphological changes are very common to many cell types involved in recruitment and migration assays and will therefore benefit a wide range of researchers in these areas.

In the following we will first describe the algorithms and techniques in image processing that are implemented by CASTLE, leading to the subsequent ImageJ plugins that implements the workflow developed in this paper. Then, we explain the methods used in evaluating the results. Finally, we gauge the performance of the designed workflow and analyse the results produced in relation to our biological application. Specifically, we evaluate the precision and recall of our automatic segmentation, and demonstrate how our high throughput analysis can be used to extract new information from data through the machine learning techniques: principal component analysis and *k*-means clustering.

## 2. Methods and Materials

In this section, the methodology of the investigation is explained. This includes details of the final automated workflow, image acquisition, and the validation process used to gauge the performance.

Images are commonly stored as a matrix with entries for each pixel. An entry’s value represent the brightness intensity. In our study we consider 8-bit grayscale images. This means each pixel varies in shade from 0 to 255, with 0 representing absolute black and 255 absolute white. The objective of image processing is to prepare an image for analysis, in our case by obtaining a binary image that identifies exactly the regions of interest (ROI). A binary image here is one where we assign pixels part of the ROI with a value of 1, and other pixels with a value of 0. For images with more than one type of ROI, multiple binary images are created and organised into channels for the respective ROI. To convert our image into the desired form, we must first pass our matrix under a series of transformations to reduce noise and to accurately identify the pixels truly part of a ROI. These algorithms of transformation broadly fit under three classes: pre-processing, segmentation and postprocessing. For each we will identify the obstacles in the acquired data; then explain the techniques used to resolve them.

Figure 1 shows a typical image used in this investigation. This image is obtained from a co-culture system where peripheral blood mononuclear cells (PBMCs) and platelets flow over an adhesive substrate. Note that the PBMCs which are initially adhered appear almost perfectly spherical and very bright, while the PBMCs in firm adhesion are comparatively dark with highly heterogeneous shape. Because of these differences, the stages of the cell are often referred to as phase bright and phase dark for initial adhesion and firm adhesion, respectively.

**Figure 1:**
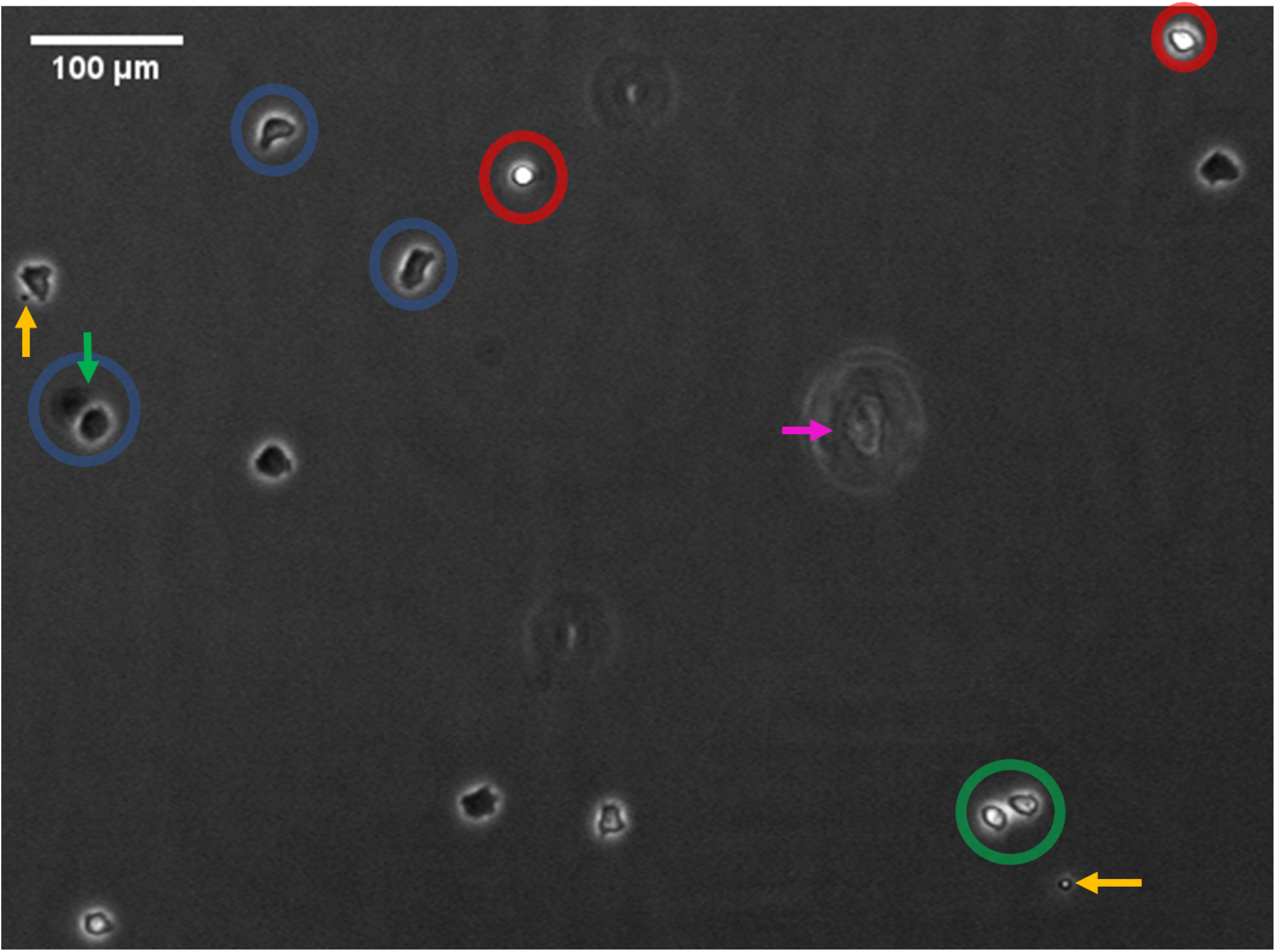
A cropped area of a representative image that can be analysed through our workflow. The image shows co-cultures with peripheral blood mononuclear cells (PBMCs) and platelets on adhesive substrates, where the PBMCs and platelets appear in different adhesive states. The image has been labelled with a number of key features identified. First, the cells we are identifying are considered to be at two different phases of adhesion, rolling adhesion (red) and firm adhesion (blue). Moreover, cells may be in a transition state between these two phases (green). All cells produce the characteristic ‘halo’ as an emanating set of bright pixels surrounding the cell. However, sometimes the ‘halo’ effect is overcompensated and can cause a following darkness in pixels (green arrow). The uneven illumination is not obvious in this image but is normally more prominent, such as in Figure 2 Irregular objects in the image are also pointed out with arrows. The orange arrows identify platelets adhered to the surface. The magenta arrow identifies a disruption in the coating of the protein causing a significant phase delay similar to a halo.

### 2.1 Pre-processing

Pre-processing uses routines to convert raw, heterogeneous images into a set of standardised images with as few unwanted artefacts and accentuate features for separation in segmentation. The techniques of PCM do introduce certain artefacts in an image which could be incorrectly identified as a ROI. During this stage we consider techniques that are used to normalise image-to-image brightness and account uneven illumination, which images like Figure 1 and Figure 2 contain.

**Figure 2:**
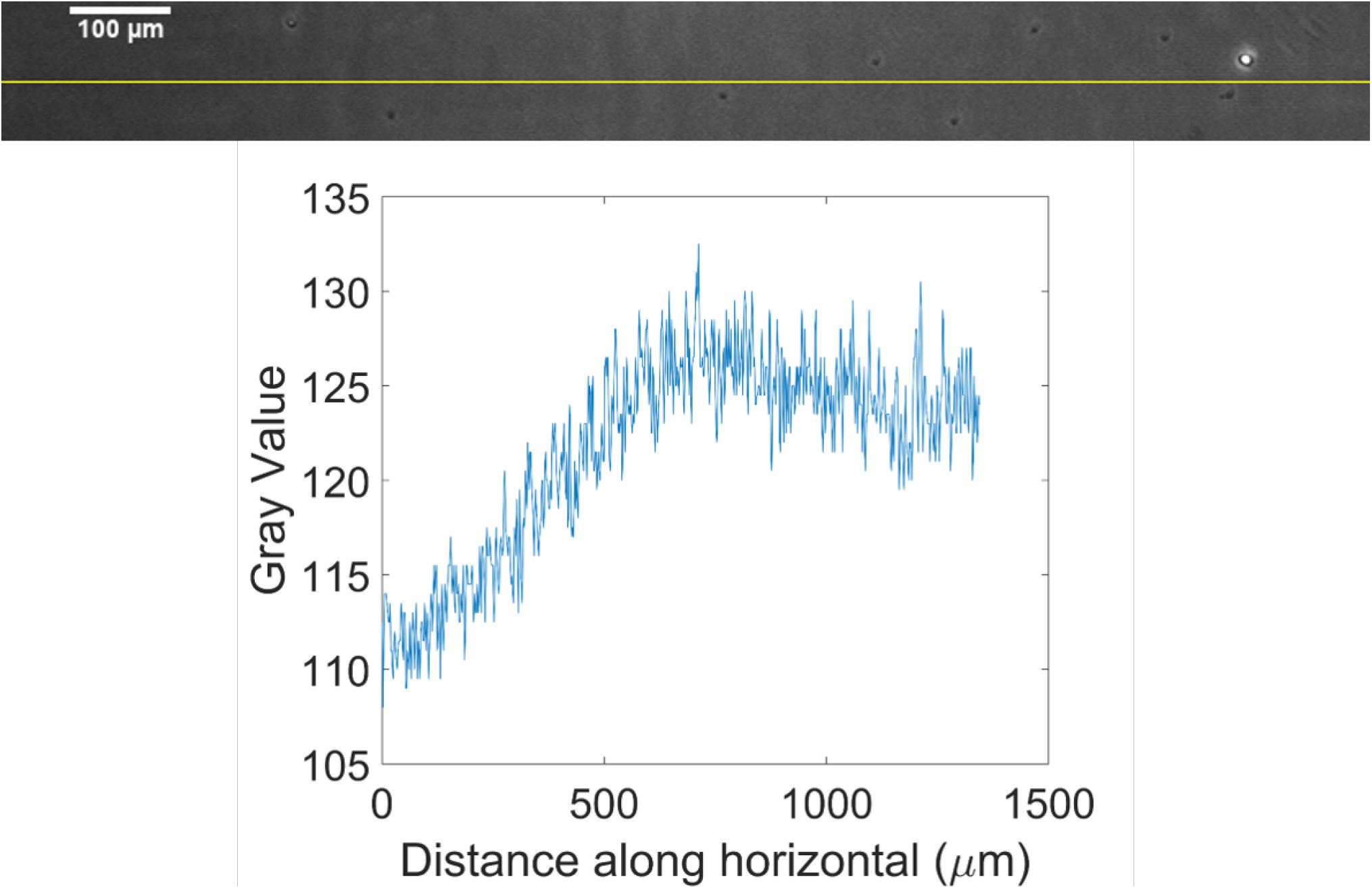
A cropped area taken from another representative image with the values of the pixels plotted against the path across the width of the image (yellow). Here we can see an uneven illumination from a light source, where the background illumination decreases in brightness the further left we go.

#### Image-to-image brightness heterogeneity

To begin eradicating the various artefacts in the image, we first normalise the pixel values so later processing techniques are repeatable across images captured with varying illumination. The average pixel intensity across a set of images, in our case sampled from the training set, and the individual difference in mean pixel intensity in each subsequent image is corrected by adding the difference to every pixel in an image so that the mean becomes that of the set. This works well for images that are considered reasonably similar. However, an image with a high proportion of ROIs which are either really bright or really dark will skew the set’s mean. This drawback could reduce the range of the pixel intensities in other images as negative values are made equal to 0 and those over 255 reduced to the maximum and so previous variations in pixel intensity are lost, which is also known as contrast.

#### Uneven image illumination

Now, we address the uneven illumination of the image and the ways to account for it. We can observe in Figure 2 that there exists an uneven illumination of the image from one side of the image to the other. We consider a technique described by Russ [8] using an averaging kernel operation with an array of large dimensions followed by image arithmetic.

The averaging filter is an iterative application of a kernel operation. The kernel operation is usually a square array of dimension (2*m* + 1) × (2*n* + 1), so that there is a central pixel, with each cell containing an integer weight. The array is put over an initial pixel at its centre. The centre and neighbouring pixels are then multiplied by the weight overlaying it. The result is summed and divided by the sum of the weights to fit the same 8-bit scale. This is then put as the value of the central pixel in the output image. This is repeated for every pixel in the input image. This can be expressed for an input image *f*, transformed to *g* by the kernel *ω* of dimension *a* × *b* as:

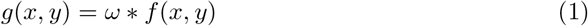

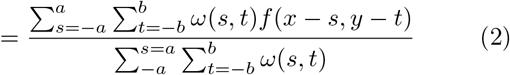

for all pixels (*x, y*) in *f*.

For pixels close to the boundary the array will extend outside the perimeter of the image. To account for this a simple method is to mirror the pixels from the boundary up to the 2*m* pixel in the horizontal and the 2*n* pixel in the vertical as padding around the image.

The techniques described by Russ is to take the average of a large area of the image, so that extreme intensities from individual ROIs do not skew the kernel operation, this is known as the *Mean* kernel. However, the area cannot be too large so that the overall differences in illumination are not lost in the output. The array of weights in *ω* need all pixels in the array to have an equal contribution, so for a 3 × 3 region we have:

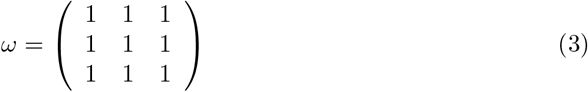

By applying a similar array with larger dimensions, fitting the properties as mentioned before, results in an output showing the changes in the background illumination. Then, the original image is divided by this background to account for the illumination across the image.

### 2.2 Segmentation

Segmentation is the identification of pixels as either associated with a ROI or not. The segmentation process works most efficiently on images which have had noise and other imperfections removed to allow for the most accurate separation of pixels to be categorised. Techniques for this stage may differ in three key areas: detection efficiency, user-operability and computational cost. The first of these relates to the success in correctly identifying the ROI. The second relates to the amount of user knowledge needed in being able to use the routine. The last analyses the time taken to execute once the program is running. These were the properties considered when designing this automated workflow. We first describe the alternative techniques considered for this methodology before describing the active learning segmentation used in the final methodology.

#### Thresholding

A key component of segmentation is thresholding. Thresholding involves an input of minimum or maximum pixel intensities which a ROI has been identified to have. If the information for setting such limits is known, then the process is simple: pixel-by-pixel, an if-statement transforms an 8-bit image to a binary image with the pixels part of the ROI as 1 and the rest 0. However, in many cases the best thresholds to choose are unknown, or they are required to be automated. There are many published algorithms that try to resolve this problem as they depend on the type of image histograms produced after pre-processing [16, 17, 18]. These have then been integrated into workflows to form some of the current techniques used for segmentation, known here as automated thresholding workflows.

#### Automated thresholding workflows

Usually a single or multiple threshold value is unable to differentiate between ROI and other artefacts that may have pixels of similar intensity. Therefore, other techniques are used to utilise common features in a set of images to aid the classification of a true ROI.

Two very common effects in phase contrast images are the characteristic ‘halo’ corresponding to the refractive index and thickness values between the cell and the surrounding medium.

The halo effect is exploited in the work of Selinummi et al. [19] and Flight et al. [20] in dense colonies of epithelial cells. The intuition behind both techniques is by applying a *Gaussian blur* or *Mean* kernel operation for a variety of different dimensions and then monitor the variation in pixel intensities. When the blur is applied, the halos are already close to maximum pixel intensity so do not vary but the darker region of the epithelial cell quickly become brighter as the surrounding halo encroaches on the ROI. Therefore, the image is transformed to one with higher pixel intensity representing higher variations. A subsequent application of Otsu thresholding [16] determines the threshold value to separate the halo, and thus background, from the ROI. This was combined with fluorescent dyes identifying constituent parts of the ROI, such as the nucleus, and Discrete Mereotopology to validate the segmented area as a ROI [21]. However, the dependency on densely packed cell colonies made these techniques unsuitable for the images captured here.

#### Active learning segmentation

Active learning is a type of machine learning method where a user iteratively adds training data to a supervised learning algorithm until the desired classification is achieved. In image analysis, a classifier is a set of rules based on active learning methods to identify the constituent parts of images not originally trained upon. The constituent parts are the different types of ROI mentioned before, and are referred to as classes in classification. The process can be applied in our workflow by creating a classifier from a random sample of the data and then using it during our segmentation step to identify the different classes of ROIs in the rest of the images.

There are many active learning software tools for segmentation already available in the bio-imaging community [3, 4]. The majority of these use an algorithm called a random decision forest as the foundation for the classifier [22]. This is whereby a multitude of decision trees are constructed during training. Then, when applied to a new object to classify, the output class is the modal class from all of the decision trees.

Ilastik is one such interactive learning and segmentation toolkit [23]. The *Pixel Classification* module calculates 35 classification features from an image matrix to interpret information about intensity, edge, texture and orientation. These help produce a classifier to be applied to other images to provide a prediction of the designated class for every pixel. Adjacent pixels of the same class then form our ROIs. So, unlike before, we are able to train the classifier to the kinds of images specifically collected with our research. In addition, the user-friendly graphical user interface (GUI) is well-suited for easy operation and limited user knowledge. Finally, there is some additional computational cost with the added complexity of the algorithm although this was still significantly faster than the current manual approach, mentioned later in the results.

### 2.3 Post-processing

Post-processing describes the type of techniques used to sieve through a binary image produced by segmentation to remove ROI that are in fact irregular particles. An example are the platelets pointed out in Figure 1. These cells could be detected as smaller versions of cells as they undergo respective stages of adhesion albeit on a smaller scale. Such techniques range from counting only those ROIs that are above a minimum and/or maximum number of pixels to machine learning algorithms separating unusual artefacts by measuring multiple morphological features. In our final workflow, a minimum and/or maximum threshold for the number of pixels in a ROI is set as the method used in post-processing. This is because the cells being identified follow fairly homogeneous measurements in size for the adhered area or are significantly distinct from particles other than that being considered.

### 2.4 Analysis

We now demonstrate the effectiveness of our approach by analysing images with adherent cells. We first quantify the number of cells depending on their adhesion status. This is done for different experimental conditions; e.g. by varying concentrations of molecules that are suspected to affect cell adhesion. Afterwards, we utilise a by-product of the automated workflow and begin to quantify the morphology of the cells at the different experimental conditions.

As the images are segmented pixel-by-pixel, to whether a pixel belongs to the ROI or not, we have a representation of the adhered surface area of each cell. As seen in Figure 1, during initial adhesion the cell remains in a highly circular form, however during firm adhesion the cell migrates on the surface looking to pass through. The shape of the cell can also be affected by surface proteins and can therefore provide useful information about the impact of molecules on the cells. Thus, an interest is to categorise the morphology of the firm adhesion cells to investigate the effect of different levels of molecules in this process. We describe the machine learning techniques used to analyse the morphology of phase dark cells.

#### Principal Component Analysis

Principal Component Analysis (PCA) is a methodology to reduce the dimensionality of a data set in which there are a large number of interrelated variables. The methodology is explained in more detail by Jolliffe [24] or Goodfellow et al. [25]. Here each ROI identified has 10 values calculated from shape descriptor formulae, refer to A.1 for exact formulae. The result is a representation whose elements have no linear correlation with each other. The order depends on the variance in that dimension. The representations, called principal components, provide a summary ordering the most influential factors in describing morphology. Here it was reasonable to not reduce the dimensionality of the data, however the technique is useful to to display the most significant contributions to the variability for a user to interpret. The relationship between variance and information is that, the larger the variance carried along the axis, the larger the dispersion of the data points along it. Therefore, if the dispersion in that dimension is larger, the particles are more likely distinguishable.

#### k-means Clustering

The *k*-means clustering algorithm divides the data set into *k* different clusters of data-points that are considered close to each other. First, each ROI is represented by a vector with each entry corresponding to the value of a feature. The algorithm works by initialising *k* different centroids to some ROI. Afterwards, the algorithm assigns each ROI to the cluster with minimal Euclidean distance in the feature space to the centroid of that cluster. Then, each centroid *μ_i_* is updated to the ROI closest to the mean of all the training examples assigned to cluster *i*. The algorithm alternates between the two steps until convergence or the maximum number of iterations is achieved. The methodology is explained in more detail by Lloyd [26].

#### Statistical testing and inference

In our investigation, we are looking to classify the morphology of the cells of firmly adhered cells depending on the different concentrations of molecules. Therefore, using the null hypothesis that groupings are independent of the molecule we are investigating, we can test the statistical significance of the outcome in the analysis.

If proven significantly different, then the shape descriptors contributing the most to the principal components from the PCA indicate the features most effected by these differences.

### 2.5. Validation

The success of classification was measured by the workflow’s precision, recall and combined score of the two as described by Powers [27]. This is similar to the validation process used by many publications of the area ([19], [28], [29]) and hence gives a comparative statistic between different automated workflows. The statistics necessary for these calculations describe all the possible outcomes for a classifier. These possibilities are:

- true positive (TP) – correctly identifies the adhered cell of interest as a ROI.
- true negative (TN) – correctly does not identify noise as a ROI.
- false positive (FP) – incorrectly identifies noise as an ROI.
- false negative (FN) – incorrectly identifies adhered cell as not a ROI.

Classification precision (*p*), or positive predictive value, indicates the fraction of objects correctly classified as a ROI out of the total number detected, and is calculated as:

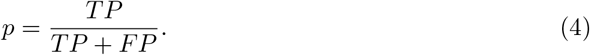

Classification recall (r), or sensitivity, indicates the fraction of objects correctly classified as a ROI out of all adhered cells, and is calculated as:

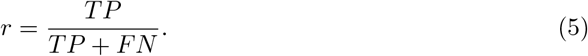

The proportions for *r* and *p* are expressed as percentages.

The *F*_1_ score is the harmonic mean of precision and recall:

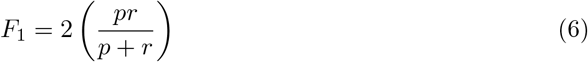

An *F*_1_ score of 0 indicates no agreement for the automated and manual analysis, while 1 indicates complete agreement. The final *F*_1_ score, precision and recall is calculated as the average of the scores from both phases.

Next we investigate where the automated workflow can be improved with image-by-image comparisons of the manual and automated number of ROI. We can categorise the incorrect detection of adhered cells into four categories: missing, noise, merged and split. Missing describes those cells that are not detected by CASTLE but counted by a human expert.

Noise are the particles identified as a ROI by the automated workflow but are not counted by the expert. Merged detections are the number of adhered cells that are counted as being part of another detected ROI. Finally, a split observation is a single adhered cell that is detected as two distinct ROIs.

### 2.6. Image acquisition

Here we describe briefly the specific experiments performed to acquire the images used to validate our workflow.

#### Blood Sample Collection

Blood samples were obtained from donations of volunteers who have given written informed consent. Approval was obtained from the University of Birmingham or Queen Mary University of London Local Ethics Review Committee.

#### Gal-9 experiments

A flow chamber assay was used to investigate the adhesion behaviour of PBMCs and purified PMNs for different amounts of Gal-9 under physiological flow conditions.

Channels in an Ibidi chamber were coated with either 20, 50 or 100 *μ*g/ml Gal-9 or 1.5 % bovine serum albumin (BSA) in Phosphate-buffered saline (PBS) as the control. During the experiment, the isolated PBMCs or PMNs were perfused through the channels at a concentration of 1 × 10^6^ in PBS containing Ca^2+^ and Mg^2+^ for 4 min before a wash out period of 1 min. A shear wall stress of 0.1 Pa was applied throughout the perfusion to mimic physiological conditions.

After the wash out period, a series of images was taken under continuous flow across the channel. The cell number adhered per mm^2^ and per 1 × 10^6^ cells was calculated from the series of images.

#### ICAM-1 experiments

A flow chamber assay was used to investigate the adhesion behaviour of purified PMNs on ICAM-1 under physiological flow conditions.

Channels in an Ibidi chamber were coated with 10 *μ*g/ml ICAM-1-Fc. During the experiment, the isolated PMNs were perfused through the channels at a concentration of 1 × 10^6^ in PBS containing Ca^2+^ and Mg^2+^. PMNs were allowed to enter the chamber and then flow was suspended for 5 mins to allow adhesion of cells to ICAM-1-Fc. Flow was then restarted at a wall shear stress of 0.1 Pa, which was applied throughout the perfusion to mimic physiological conditions.

After the wash out period, a series of images was taken under continuous flow across the channel. The total cell number adhered was calculated from the series of images.

#### vWF experiments

A flow chamber assay was used to measure adhesion of platelets to von Willebrand Factor (vWF) under physiological flow conditions.

Glass microslides were coated with 0.1 mg/ml vWF and blocked using PBS containing 2% BSA. During the experiment, a flow rate of 0.8 ml/min was maintained to give the desired wall shear stress of 0.1 Pa. Anti-coagulated human whole blood was perfused for 2 minutes over vWF. The microslides were then washed with PBS 0.1% BSA with or without 30 *μ*mol/l Adenosine Diphosphate (ADP), a platelet activating agent.

After the wash out period, a series of images and videos were taken under continuous flow across the microslide. Platelet coverage was calculated as the number of cells adhered. Phase bright platelets were classed as initially adhered platelets and phase dark platelets were classed as activated platelets.

### 2.7. Image processing and analysis using CASTLE

The automated workflow is visualised in Figure S1. The batch processing was designed as a script in the ImageJ Macro language and implemented as a plugin in Fiji is Just ImageJ 2.0.0 [30]. The *Mean* and *Image Calculator* plugins were used in Pre-processing [31]. Training of the classifier and Segmentation was carried out using Ilastik 1.3.2post2 [23]. Post-processing and data collection for analysis were completed using the *Analyse Particles…* plugin [31]. Analysis was carried out in R 3.6.1. [32]. The final automated workflow is available as a collection of open-source plugins on the Cell Adhesion with Supervised Training and Learning Environment (CASTLE) GitLab project, refer to Section 5. The computer used in the implementation of the CASTLE plugins had an Intel(R) Core(TM) i7-9850H CPU at 2.60GHz with 16GB RAM.

## 3. Results

We now apply our workflow to several data sets. We focus most of our discussion to the analysis of PMNs and PBMCs depending on concentrations of Gal-9, since these data sets are the most challenging to the algorithms and involve different cell types with quantitatively varying concentrations of molecules. We then demonstrate that our workflow can similarly be applied to the same cell types stimulated with other molecules (here, ICAM-1), or very different cell types (here, platelets).

### 3.1. Validation of workflow

#### Validation of cell count

The automated workflow, CASTLE, was applied to the images from an ongoing study of the role of the protein Gal-9 on leukocyte adhesion. For each of the 5 different levels of protein exposure, we obtained a sample of 7 different images. One image from each protein exposure was chosen for the classifier to be trained upon. After the automated workflow was applied, the plugin took under 11 minutes for all 35 images to be processed (see Table S1). One image was excluded from the validation after statistical analysis identified the image as an anomaly compared to similar images of the same protein exposure (see Figure S2). The final *F*_1_ score was 87.0%, with a 81.2% classification precision and an 93.7% classification recall.

The final validation scores were the result of introducing a third ROI for classification. This was identified in Figure 1 as a cell transitioning from initial adhesion to firm adhesion, which in manual counts were associated with the latter but were considered still too bright in the automated workflow. Before the new class was introduced, the *F*_1_ score was 86.2% with a classification precision of 89.9% and a classification recall of 82.8%.

Besides the increase in *F*_1_ score for the count, the introduction of the new class was preferred as the following analysis into morphology depended on correctly identifying all of the cell that forms its shape. Using the same classifiers, a validation on all the classified pixels was carried out on a sample of 5 images, one from each of the different cell type and protein exposures. The 5 images were segmented manually by an expert to identify the phase dark cells. The same 5 images were processed by CASTLE. The images were then compared pixel-by-pixel. Before the introduction of the Transition class during training, the *F*_1_ score was 69.6% with a classification precision of 85.0% and a classification recall of 59.0%. The *F*_1_ score remained similar at 69.3% but with a higher classification precision of 89.2%, the outcome preferred, and a classification recall of 56.7%.

Figure 3 shows the analysis carried out on the performance of the automated workflow compared to that counted by an expert. A perfect identification of the number of cells would result in the percentage of correctly identified cells in the respective phases equal to 100% of the true count. We see across the phases of cells identified that the majority of incorrect counts are due to noise and missed cells. Noise adds to the total count by leading to false positives, while missed cells reduce the final count. Figure 3b confirms that the introduction of a class for transitioning cells significantly reduces the number of incorrectly classified cells (i.e. improving the precision) and the number of cells missed (i.e improving the recall) compared to the classification based on two classes in Figure 3a.

**Figure 3:**
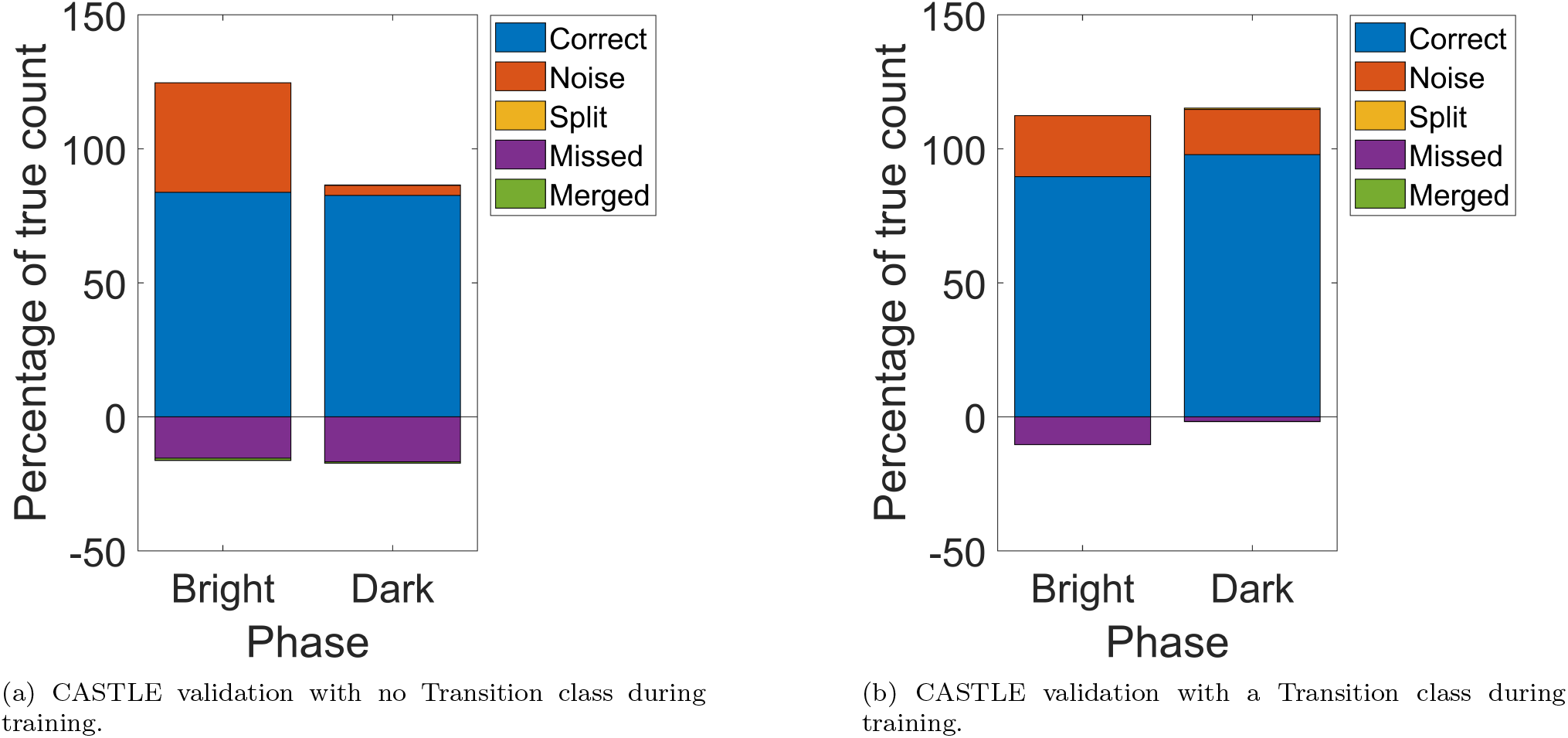
Validation of the automated workflow against manually counted and verified results from the study into the role of the protein Gal-9 on leukocyte adhesion. (a) CASTLE validation with no Transition class during training. (b) CASTLE validation with a Transition class during training. One image from both analyses was excluded after being identified as anomalous. The stacked bar chart shows, of all the manually counted cells, how the automated workflow performed, including how mis-identified cells of the automated workflow effected performance as a percentage of the true number of cells.

Similar analysis for the validation of CASTLE was carried out on the other studies, one on leukocytes exposed to the protein ICAM-1 and another on platelets exposed to the protein vWF. The final *F*_1_ score was 87.0% and 80.7%, respectively. Similar observations were made to that already discussed (see Tables S2, S3 and Figures S3, S4). This time the iterations of training were carried out to demonstrate the application of CASTLE. First, we validated how CASTLE processed the images using the final training used in the study of the role of the protein Gal-9 on leukocyte adhesion. Then, we validated how CASTLE processed the images with an additional image from the new data set, respectively. The rationale was to show how CASTLE may be applied to new data sets: by adding to the original data set of a similar study. The result is a more thorough training due to more images identifying the variability in cell appearance yet not at the expense of the researcher’s time, who would have otherwise trained the data themselves.

#### Comparison to other methods

To confirm that the performance of CASTLE was an improvement to other available workflows, we compared the software to a comparable plugin in ImageJ/Fiji. The plugin is called Trainable WEKA Segmentation (TWS) [33] and forms part of the Waikato Environment for Knowledge Analysis (WEKA) collection for data mining with open-source machine learning tools [34]. Using the same pre-processing and post-processing techniques as CASTLE on the same 35 images, TWS produced a final *F*_1_ score of 63% with 48% precision and 95% recall (see Table S1). TWS identified a similar number of correct cells as CASTLE. However, for this data set, TWS seemed to mis-identify as many objects which increased the final number detected by about twice as many for both phases of cell (see Figure S5). Also, the time to run TWS to batch process all 35 images took 6 times longer than the comparable application by CASTLE (see Table S1). Therefore, the CASTLE workflow would appear to be an improvement in accuracy and speed than what is currently available.

#### Distinguishing cells types and experimental conditions

In addition to the total count across all of the images, an investigation into the performance of the program for the full range of cells in a particular image is presented in Figures 4 and 5. These plots display the number of cells counted manually against those detected by CASTLE. A line of least square regression is drawn to predict the program’s performance across the range of cell numbers in comparison to another line for what would be considered a perfect count, where the detected number of cells equal those identified by the expert.

**Figure 4:**
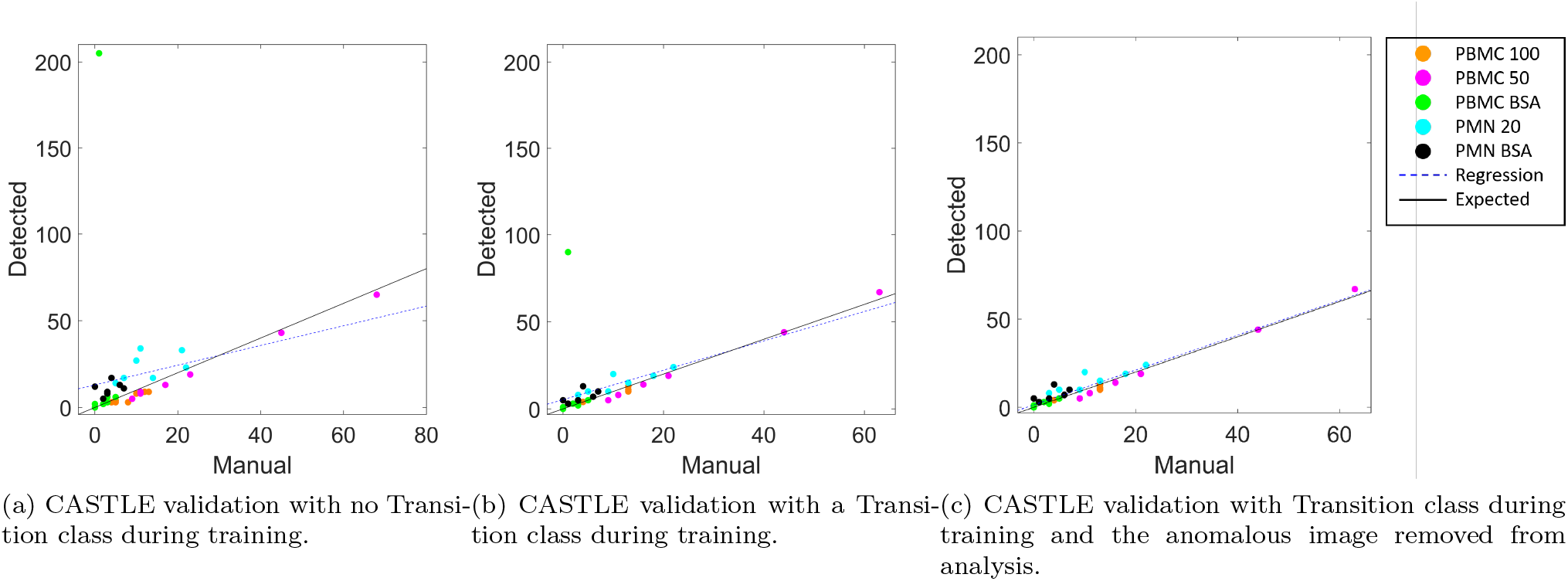
Comparison of automatic to manual detection of adherent phase bright cells for each image in the study into the role of the protein Gal-9 on leukocyte adhesion. (a) CASTLE validation with no Transition class during training. (b) CASTLE validation with a Transition class during training. (c) CASTLE validation with Transition class during training and the anomalous image removed from analysis. The regression line (blue, dashed) shows a prediction for the detected counts across the range of cells in any one image. The line of perfect agreement (black, solid) is the desired outcome with the manual count equal to the automatic count.

**Figure 5:**
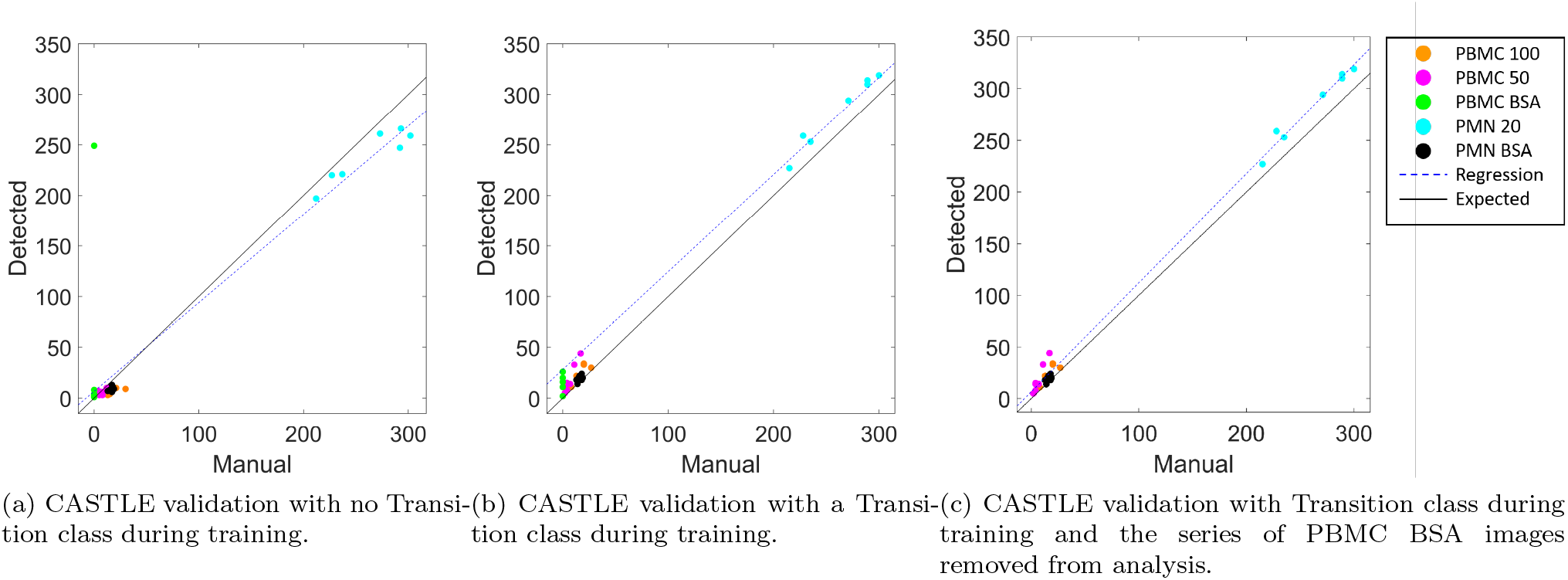
Comparison of automatic to manual detection of adherent phase dark cells for each image in the study into the role of the protein Gal-9 on leukocyte adhesion. (a) CASTLE validation with no Transition class during training. (b) CASTLE validation with a Transition class during training. (c) CASTLE validation with Transition class during training and the series of PBMC BSA images removed from analysis. The regression line (blue, dashed) shows a prediction for the detected counts across the range of cells in any one image. The line of perfect agreement (black, solid) is the desired outcome with the manual count equal to the automatic count.

Figure 4 is the series of least regression plots when comparing the performance of CASTLE to a manual analysis when confronted with the images of our study for phase bright leukocytes. Figure 4a is the phase bright analysis for all of the images when CASTLE is trained without a Transition class. However, the performance of the program was seen to still be detecting more cells which would be considered as transitioning from phase bright to phase dark, which an expert would consider as phase dark in their counts. After introducing a Transition class in the training, we see in Figure 4b an improvement in the accuracy reflected by the closer alignment of the regression line to the line representing a perfect count. Nonetheless, we can see that an image with PBMCs exposed to the control protein BSA has a significantly larger number of cells detected in comparison to the other 6 images of the same experiment. On investigation of this group of images, an image appeared to have many irregular particles seemingly trapped under the surface of the protein which induced enough of a phase difference to create the ‘halo’ effect of adhered cells (see Figure S2). After excluding the anomalous images from the input data we produce the results in Figure 4c. Therefore, with the introduction of a new training class and the exclusion of the anomalous image the final performance across the different cell types and their respective protein exposures produced a significant improvement in the correct number of detected cells.

Figure 5 is the series of least regression plot for comparing the performance of CASTLE to a manual analysis when confronted with images from our study for phase dark leukocytes. Figure 5a shows the performance when both the Transition class is yet to be introduced to the training and before the images of PBMCs exposed to BSA were analysed for anomalies. The regression line indicates that CASTLE was for the majority of images accurate but diverges to underestimate the counts for ever larger numbers of cells. Then, the Transition class was introduced into the training and the performance across the images is shown in Figure 5b. Here we are unable to observe the point representing the anomalous image mentioned before as the 521 detected exceeds the limits of the axes, the axes chosen to focus on comparing regression lines between each stage (for the inclusion of the anomalous image, see Figure S6). The regression line here shows the trend of CASTLE consistently overestimating the number of detected cells. In addition to identifying the anomalous image, after further investigation, all the images with PBMCs exposed to the protein BSA were found to have no phase dark cells and so were subsequently excluded from further analysis using phase dark cells. Using this knowledge, Figure 5c shows the final comparison after excluding these images. We know from our earlier validation that this final iteration is an improvement in *F*_1_ score from the first. For larger numbers of adhered cells in an image, the regression line is above rather than below the line of perfect count. However, in the final iteration, larger values detected do not exceed more than 14% of the manual count, unlike the 16% from before, and thus represent a lower proportion of additions and reason for improvement from the first iteration.

### 3.2. Effect of Galectin-9 on Leukocyte Adhesion

We now demonstrate how our workflow can lead to new biological insights by further analysis of the segmented images from the data set investigating the effect of Gal-9 on leukocyte adhesion.

#### Count

The purpose of the data that we analysed with CASTLE was to determine the effects of Gal-9 on cell adhesion. We now present the analysis performed by CASTLE. This is to demonstrate the ability of the program to process images into information easily interpreted by a researcher. The result is a comparison between the protein exposures within each cell type. This was carried out in the form of a Student’s t-test, where the sample variance was assumed to be unequal for each series of images. This was an assumption made in the analysis and any future use of the R script provided on the GitLab project (see Section 5) allows the use of images assumed to have an equal variance.

Our primary goal was to investigate the role of Gal-9 on the adhesion of PBMCs and PMNs. Table S4 displays the results from the Student’s t-test for each pairwise comparison of protein exposures between similar cells. At the 99% confidence level, we can conclude that for PBMCs there is a statistically significant effect on adhesion when a surface is coated with either 50 or 100*μ*g/ml of Gal-9 compared to the control. There is no statistically significant difference in adhesion between 50 or 100*μ*g/ml Gal-9 applied, at either the 99% or 95% confidence level. For PMN, the difference in cell adhesion between 20*μ*g/ml of Gal-9 compared to the control is considered statistically significant at the 99% confidence level. Figure 6 is a visual representation of these differences between protein exposure and their effect on the number of adhered cells per mm^2^ per 10^6^ cells flown through the chamber.

**Figure 6:**
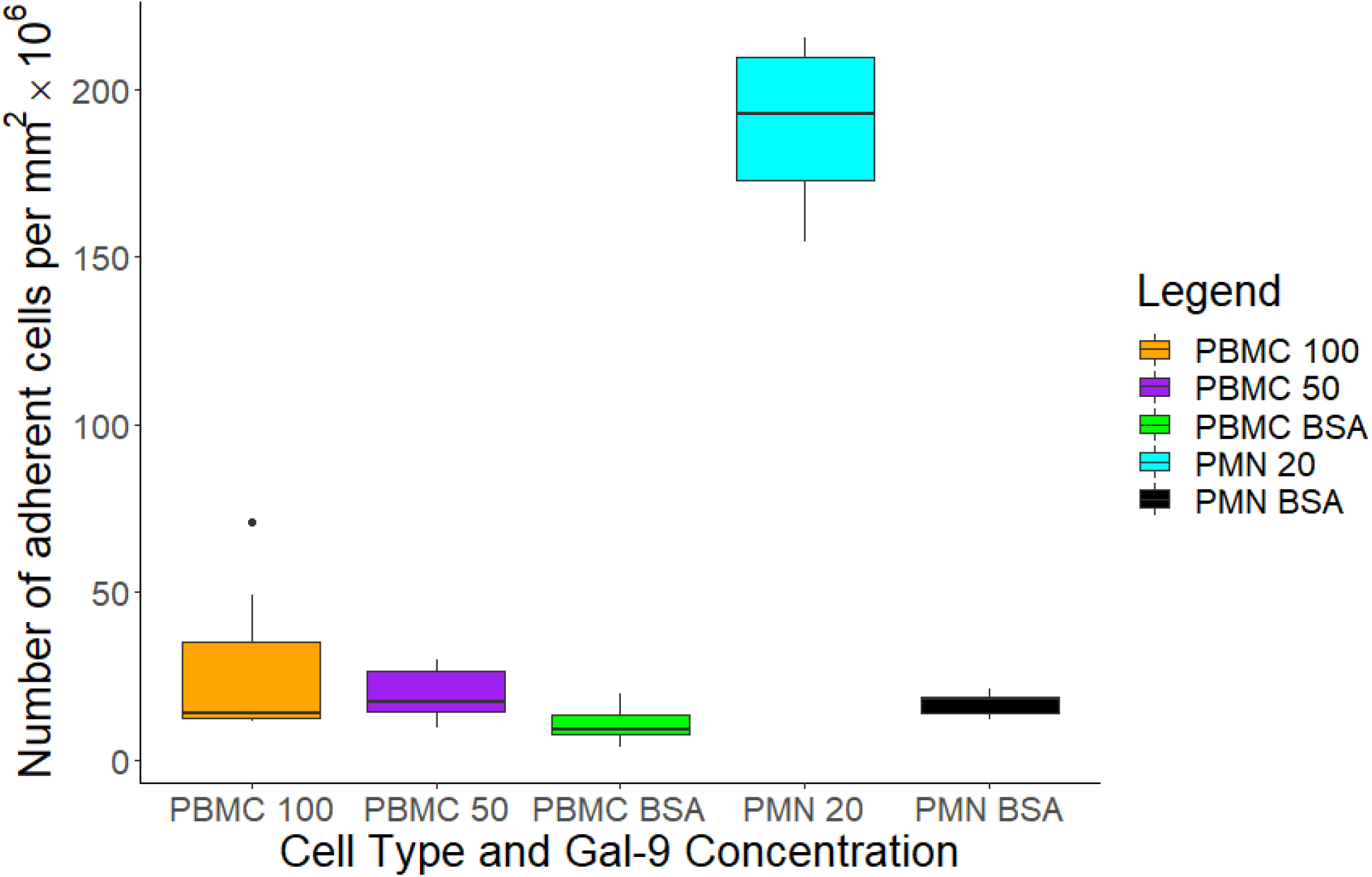
Boxplot for the total number of cells adhered per mm^2^ × 10^6^ from the sample of seven images taken for each protein exposure. This excludes the previously discovered anomalous image from the PBMC BSA group (see Figure S2).

A secondary outcome of the research was to discern whether more cells were remaining in a state of initial adhesion or whether they would form a firm adhesion in the period between perfusion and when the images were taken.

First, we consider Table S5 which displays the results of the Student’s t-test for cells during initial adhesion, known as phase bright cells. We are able to conclude, at the 95% confidence level, that there is a statistically significant difference between the mean number of phase bright cells when exposed to Gal-9 compared to the control. However, the difference at either the 99% or 95% confidence level is not considered statistically significant for distinct levels of Gal-9. For PMN, the difference in phase bright occurrence for 20*μ*g/ml of Gal-9 compared to the control is also not considered statistically significant at the 99% or 95% confidence levels.

Second, we compare the number of firmly adhered cells, known as phase dark cells, between similar cells that have been exposed to the different levels of Gal-9. Table S6 displays the results of the Student’s t-test for each combination of pairwise comparisons. For PBMCs, we can conclude at the 99% confidence level that there is a statistically significant difference in the number of adhered cells that are phase dark when Gal-9 is present on the surface at both levels. On the other hand, the number of phase dark cells between the two amounts of Gal-9 were considered not to be statistically significant at either the 99% or 95% confidence level. For PMNs, the number of adhered cells that were phase dark when 20 *μ*g/ml Gal-9 was applied, is considered statistically significant at the 99% confidence level.

Figure S7 allows us to visualise the differences in the total number of adhered cells detected for each protein exposure, with the number of phase bright and phase dark cells that contribute to the total shown. Here we can see the different effects that Gal-9 appears to have on the differing cell types. For PBMCs, the Gal-9 appears to have the largest effect on the number of phase bright cells detected in the image compared to PMNs having an enormous increase in the number of phase dark cells.

#### Morphology

The final analysis for the series of images is the shape of the adhered area between the cell and the surface coated in the protein. The following are the results from this analysis.

Table 1 displays the categorisation of the cell morphology across all protein exposures for each type of cell. First, observe that the series of images taken for PBMC cells exposed to the control protein are not present. Similar to before, after investigating the anomalous image it was seen that the corresponding series of images contained no phase dark cells. Therefore, none are in this analysis. Next, the *k*-means clustering was performed on 4 groups and so *k* = 4 in this analysis. Consider the row reflecting on the categorisation of phase dark PMN cells exposed to 20*μ*g/ml of Gal-9 to observe how well the *k*-means clustering has been able to separate the cells depending on their morphology. We see that out of the phase dark leukocytes detected, the majority are split evenly between the 2nd, 3rd and 4th cluster. This suggests that the *k*-means clustering was unable to differentiate the morphology compared to the other cells and protein exposures. However, the majority of the detected PMN cells exposed to the control were also grouped into the 4th cluster. This suggests that there may be similarities within each of the different cell types which are interfering with the heterogeneity between protein exposures. Two more *k*-means clustering analyses were carried out considering only the same type of phase dark cells.

**Table 1:** Results of the *k*-means clustering on all 2496 phase dark cells detected by CASTLE. The values represent the proportion of cells compared to all phase dark cells detected. We then compare to the number of each cell type and their exposure to the subsequent clusters that cells with similar morphological features have.

|  | Cluster 1 | Cluster 2 | Cluster 3 | Cluster 4 |
| --- | --- | --- | --- | --- |
| PBMC 50 | 0.0280 | 0.0021 | 0.0042 | 0.0102 |
| PBMC 100 | 0.0285 | 0.0063 | 0.0080 | 0.0285 |
| PMN 20 | 0.0787 | 0.2127 | 0.2876 | 0.2540 |
| PMN BSA | 0.0165 | 0.00127 | 0.0119 | 0.0208 |

Table 2 shows the analysis in morphology of the PBMC cells exposed to the different levels of Gal-9. The first observation is the relatively even distribution of cells between both clusters for each level of protein exposure. Thus, there appears to be a difference in morphology within the class of phase dark cells which is more apparent than the difference caused by the adhesion to Gal-9.

**Table 2:** Results of the *k*-means clustering on the 298 phase dark PBMCs detected by CASTLE. The values represent the proportion of cells compared to the number of phase dark PBMCs detected. We then compare to the number of each cell type and their exposure to the subsequent clusters that cells with similar morphological features have.

|  | Cluster 1 | Cluster 2 |
| --- | --- | --- |
| PBMC 50 | 0.1611 | 0.2234 |
| PBMC 100 | 0.3699 | 0.2454 |

Table 3 is the result in the *k*-means clustering for phase dark PMNs. It appears that PMN cells exposed to 20*μ*g/ml of Gal-9 is still evenly distributed between the two clusters although those exposed to Gal-9 are more similar to the second cluster than the first. However, no significant distinction can be made from this analysis.

**Table 3:** Results of the *k*-means clustering on only the 2112 phase dark PMNs detected by CASTLE. The values represent the proportion of cells compared to the number of phase dark PMN cells detected. We then compare to the number of each cell type and their exposure to the subsequent clusters that cells with similar morphological features have.

|  | Cluster 1 | Cluster 2 |
| --- | --- | --- |
| PMN 20 | 0.3806 | 0.5486 |
| PMN BSA | 0.0341 | 0.0365 |

In all of the *k*-means clustering analyses of morphology we have been unable to determine a separation of a cell’s morphology depending on their protein exposure. To better understand this, we show the first two principal components in Figure 7. This shows the first two independent dimensions that capture the highest variance to help separate the data and feed into the *k*-means clustering algorithm. We choose the first two components only as they can be easily displayed. For the first analysis, shown here as Figure 7a, the PCA is not able to separate the data sufficiently enough to be easily distinguished into the different groups of cells. This seems similar to the Figures 7b and 7c for the PCA for PBMC and PMN cells, respectively. However, these plots are still able to convey relationships between the morphology and the protein exposure. For example, Figure 7a can begin to identify characteristics between the two types of cell through their morphology. Here, we are able to see in the first principal component that PBMCs appear with a negative value in the first principal component for all but a few points. This is in contrast to PMNs which the majority exposed to Gal-9 appear above 0 and those exposed to the control appearing across the full range. When considering the second principal component, there appears to be 2 general sub-classifications forming with a cluster of points centred about (1,−1) and the rest sparsely centred about (−4,0).

**Figure 7:**
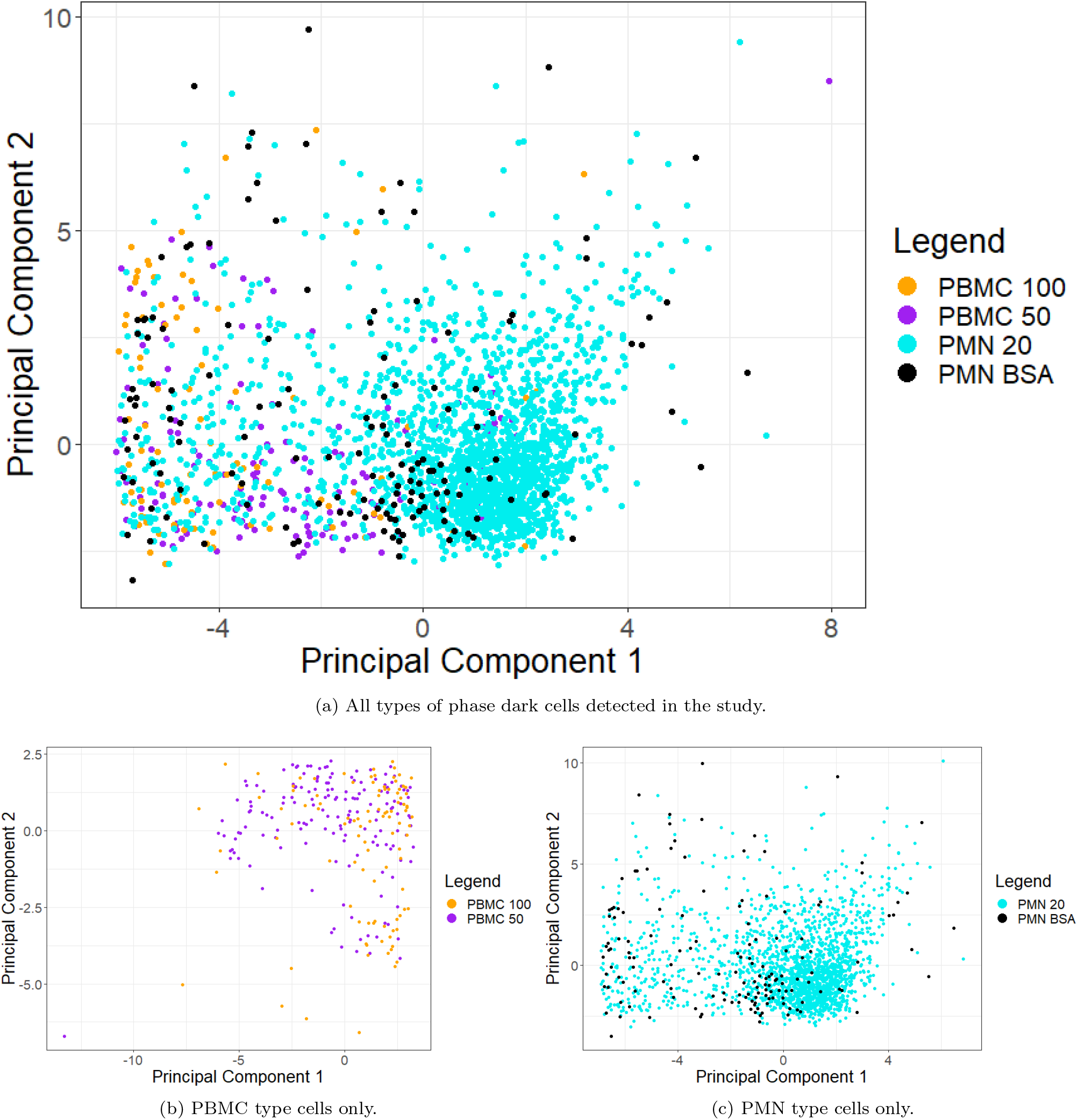
Morphology of adhered cells considered phase dark by CASTLE plotted in the first and second principal components for each of the PCAs. (a) All types of phase dark cells detected in the study. The first two principal components capture 73.27% of the total variation in cell morphology. (b) PBMC type cells only. The first two principal components capture 76.39% of the total variation in PBMC morphology. (c) PMN type cells only. The first two principal components capture 72.13% of the total variation in PMN morphology.

Figure 7b is a PCA on PBMCs only. We observe that although both groups of Gal-9 exposure have values distributed across the range of first and second principal component values, that there could be some trend. This trend is in reference to many of the PBMCs with 50 *μ*g/ml are more apparent from just lower that −5 to 2.5 in the first principal component but for mainly positive values in the second component. This is compared to the more apparent regimes for PBMCs appearing between 0 to 2.5 in the first principal component and from lower than −5 to 2.5. This is a result not obvious in Figure 7a but is more obvious with the individual analysis, most likely due to less cell heterogeneity.

In Figure 7c, inferences in trend can be made about the differences from exposure to Gal-9 to the control for PMNs. Many of the points are dispersed closely about (0,0). However, the majority of points appear for positive values in the first principal component for those exposed to Gal-9 than those that do not. On the other hand, those exposed to the control, although some begin to cluster about (−1,−1), have many values at the extremes of one or both of the principal components. Referring back to Figure 7a, this appears to be the case too. This observation could suggest that instead of exposure to Gal-9 causing new or more extreme phenotypes in the cell that it encourages a specific morphology already possible by the cells under normal adhesion. To understand what morphology a cluster represents, referring back to the shape descriptors that contribute the most to the first and second components can identify which are less variable to exposures of Gal-9.

### 3.3. Application of CASTLE to other data sets

We now demonstrate the applicability of our workflow to two other studies of cell adhesion: one investigating the effect of ICAM-1 on PMN adhesion, and the other investigating the adhesion of platelets in dependence on vWF and ADP. The biological interpretation of these results are not discussed here, but the results from CASTLE are reproduced in the Supplementary Materials to show the applicability of CASTLE to different data sets. Using a similar flow assay as for the studies into Gal-9 (see Section 2.6), we quantify the number of adherent cells in the images from both studies.

In the study introducing ICAM-1 exposure on PMNs, each image was from a repeated flow assay experiment. The results produced by CASTLE are displayed in Figure S8, showing the total number of adhered cells and the proportions of which are phase bright and phase dark.

In the study on platelets, an investigation into the role of vWF in the adhesion of platelets is considered, in dependence of ADP. As in the application of CASTLE to the first study, Figure S9 shows the total number of adhered cells and the proportions of which are phase bright and phase dark. In addition, as there was a dependence on ADP between images, Figure S10 shows the number of adherent platelets obtained at different time points within the same experiment with and without ADP. We observe a higher number of adherent cells in the presence of ADP. Moreover, Table S7 shows that the difference in total, phase bright and phase dark adhered platelets when ADP was introduced is statistically significant at the 99% confidence level. This is expected, since ADP is known to activate platelets, allowing them to bind to vWF [13, 14].

## 4. Discussion

We have introduced a new workflow for the automatic analysis of PCM images of adherent cells. We demonstrated that the performance of CASTLE is comparable to methods already established in the detection of cells undergoing similar processes. One such example are the methods used by Huh et al. [29] in the detection of mitosis in PCM images for stem cell populations. In their study, the method at greatest accuracy achieved an average *F*_1_ score of 88.6% with a 91.0% precision and 86.7% recall on a population of stem cells. Therefore a final *F*_1_ score of 87.0% with a 81.2% precision and an 93.8% recall describes a system ready for use in future research, if not a benchmark to improve upon.

The most successful property of the program is the reduction in manual interaction. For a similar analysis of the 35 images counted in the the Gal-9 study, the process takes approximately 1.5 hours to count and analyse by hand. In comparison, CASTLE is recorded taking under 12 minutes on a conventional laptop computer. Moreover, CASTLE provides tools for the analysis of shape, yielding additional information beyond simple cell counts.

CASTLE was able to give a succinct set of results for quantifying the number and proportion of cells adhered to a surface exposed to varying levels of molecules compared to a control. The CASTLE workflow determined a statistically significant difference in the number of adhered cells for both PBMCs and PMN cells depending on Gal-9 presented on the chamber surface. In addition, the case study was able to reference for which phase of cell the protein had the most impact on cell adhesion. By distinguishing between the number of phase bright and phase dark adhered cells, inferences can be made on cell exposure to Gal-9. However, the program currently has a reduced statistical power due to the accuracy of cells currently detected, demonstrated by the lower classification precision. Methods have already been considered to improve CASTLE in this aspect. One such method would be to use a series of pixel classifiers and include their output as an additional channel of information in a final pixel classifier. The intuition is called Auto-context and the core algorithm was introduced by Tu and Bai [35]. In addition, the validation process has proven that further understanding of the problem at hand and refining the workflow produces improved results. The final iteration of least square regression plots, Figure 4c and Figure 5c, are prime examples for where the line of regression between each previous iteration improves with each resolution of the processing problems. First, a new class was introduced in training, with the majority of spurious ROI eradicated. Then, the anomalous images were excluded and an improvement in the final performance of CASTLE.

The pixel-by-pixel output by CASTLE was lower in classification precision and recall than the counting performance. This is to be expected as detection holds a lower threshold than the additional identification of each pixel within that cell. This can be seen as one of the inaccuracies in the subsequent morphology analysis. Without a higher *F*_1_ score the statistical power of the *k*-means clustering remains lower. Likewise, the aforementioned Auto-context process would be able to improve the identification of pixels belonging to a cell in addition to detecting a cell as a ROI. The morphology of the phase dark cell still gives an insight into the effect of Gal-9 on the cell’s migration across the surface. Despite the methods used being unable to distinguish differences between the morphology, the workflow was able to efficiently process each cell and the 10 features describing their morphology into clusters to show their similarity clearly to a user, with the first principal component in the PCA describing the most influential factors in that separation. Without the use of the machine learning algorithms both computation and interpretation of the 10 dimensional data set would be too inefficient to process and the information forfeited. Nonetheless, limitations in the assessment of morphology are apparent.

The shape descriptors used as features could be considered insufficient for describing the complex and heterogeneous shape displayed by the migrating cell. While *Area* and *Perimeter* are tangible measures of shape, the remaining features are approximations of how similar the appearance is to well-defined shapes. The measures can become problematic when the indicator has a result at neither extreme: either the region in question is almost exactly the same as the comparable shape or not at all. When this occurs a small difference in the calculation can indicate vastly different outcomes in the true shape. Examples comparing and contrasting different dimensionless statistics of shape are described by Russ and Russ [36]. To avoid misconceptions on shape, either the features chosen should be verified by the user, then exposing the analysis to human bias, or more interpretations of shape need to be considered to lessen the weight of other measures, thus reducing the chance of peculiar similarities between heterogeneous cell shapes.

An alternative approach is to rethink the current morphological analysis and integrate previously validated machine learning techniques from similar fields. A candidate for this proposal is the work of Yin et al. [37] where a Support Vector Machine (SVM), a supervised machine learning model, was used to identify discrete shapes from 211 geometric and texture properties of each segmented *Drosphilia* Kc cell. Another example is the work of Wu et al. [38] with their image analysis tool VAMPIRE. The tool reduces the morphological description for an outline into similar modes of shape using PCA before analysing their distribution in cancer metastasis. This work also introduces analysis of cell-cell contact, possibly relevant to the aggregation of platelets. Finally, a work that does not depend on machine learning, or even human intervention, is that of Sanchez et al. [39]. Here, they develop a method called lobe contribution elliptical Fourier analysis (LOCO-EFA), an extension of elliptical Fourier analysis, whereby a contour is reconstructed from an infinite summation of related ellipses. The summation of ellipses then infer information about the original shape of the cell, in this case complex shaped pavement cells of Arabidopsis thaliana wild-type and speechless leaves, and Drosophila amnioserosa cells. Future extensions of the automated workflow could consider incorporating these morphological analysis tools.

Alongside the quantification of still images, the workflow used in CASTLE can be easily applied to videos. Videos are characterised by computers as a series of still images with a time dependence. Hence, the successful classification of multiple images is similar to a classification of a whole video with an ordering. Combined with other programs in cell tracking, such as Mosaic Suite’s Particle Tracker [40], the work can be extended to observe the whole duration of adhesion for a particular cell. Thus, other features like migration tracking can be investigated.

The methodology behind CASTLE is not limited to cell classification. The Ilastik program allows for the possibility of other types of images to be segmented into any number of classes. The utilisation of the automated workflow is only limited by the universality of pre- and post-processing techniques. The larger impact of the work is yet to be tested in other areas, such as introducing a monolayer of endothelial cells to monitor extravasation.

Here we have designed an automated workflow, from data acquisition to analysis, which can be applied to numerous PCM images. A fundamental understanding of the biological processes allowed for the method to be refined to obtain an accurate conclusion. It was shown that the results were comparable to the manual analysis of the same data set and produced information beyond what was previously available. In addition, these conclusions can be reached significantly faster than the alternative manual method with minimal expense to a researcher’s time. Potential ways to enhance the performance are already being investigated and new lines of interest are being considered for application.

While much of our discussions focused on the investigation of adhesion of PMNs and PBMCs in dependence on Gal-9, we demonstrate the applicability of our workflow to other cell types and other experimental conditions. Notably, we investigated lecukocyte adhesion in dependence on ICAM-1, and platelet adhesion to vWF in dependence on ADP. Strikingly, cells were well recognised even when using the training set from the Gal-9 studies. Recognition can then be improved by adding single annotated images to the training set, minimising the manual input for this automatic analysis. This demonstrates the utility of CASTLE for the analysis of a wide range of adherent cells under various conditions.

## 5. Software availability

The ImageJ macro scripts created for the purposes of this research, namely Cell Adhesion with Supervised Training and Learning Environment (CASTLE), are available to download from the following project on GitLab: https://gitlab.bham.ac.uk/spillf/castle. The R script to perform the Principal component analysis (PCA) and *k*-means clustering algorithms, to analyse the results from CASTLE, are available at the same location. Finally, the validation performed is also available as a series of spreadsheets at the same location.

## Acknowledgments

The authors would like to thank Jeremy Pike and Iain Styles for discussions and advice during the preparation of this article. SGG is supported in part by the EPSRC Centre for Doctoral Training in Topological Design, funded by the UK Engineering and Physical Sciences Research Council (grant EP/S02297X/1) and the University of Birmingham. MC was supported by a Royal Society Dorothy Hodgkin fellowship (DH160044). AJI is supported by the University of Birmingham Fellowship and AMS Springboard Award (SBF003/1156). The Institute of Cardiovascular Sciences, University of Birmingham is recipient of a BHF Accelerator Award (AA/18/2/34218).

## A. Supplementary Material

### A.1 Shape Descriptors

The following are morphological features available as measurements for shape analysis in the image processing software ImageJ as defined in the user guide [31].

#### Area

Number of square pixels contained in a ROI, or in calibrated square units (e.g., mm^2^, *μ*m^2^, etc.).

#### Perimeter

The length of the outside boundary of the ROI.

#### Bounding rectangle

The smallest width and height of the rectangle enclosing the ROI.

#### Fit ellipse

Fits an ellipse to the ROI. Parameters Major axis and Minor axis are the primary and secondary axis of the best fitting ellipse.

#### Circularity

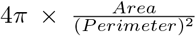 with a value of 1.0 indicating a perfect circle. As the value approaches 0.0, it indicates an increasingly elongated shape.

#### Aspect ratio

The aspect ratio of the particle’s fitted ellipse, i.e., 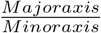.

#### Roundness

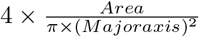

#### Feret’s diameter

The longest distance between any two points along the selection boundary, also known as the maximum caliper. In addition, the minimum caliper diameter is calculated, also known as the minimum Feret’s diameter.

---

**Figure S1:**
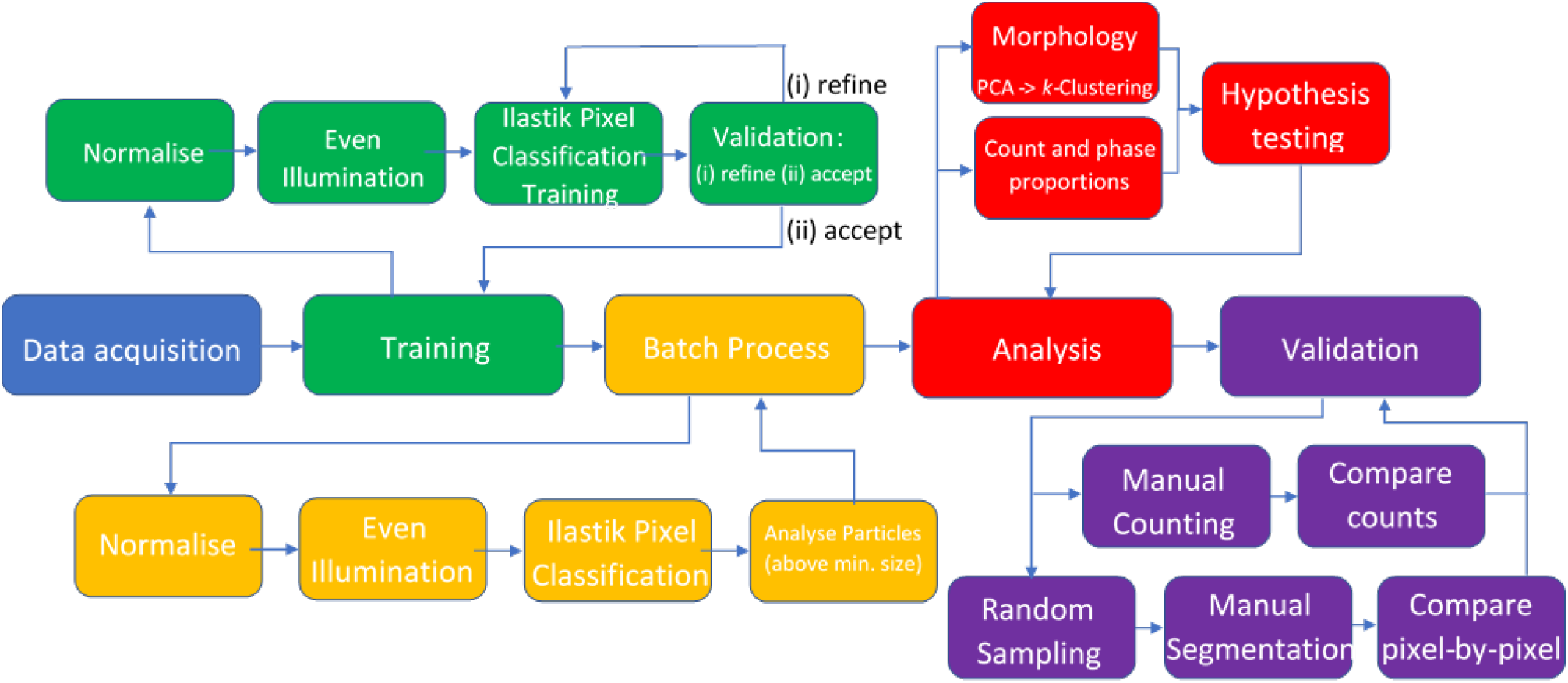
The workflow for the application and validation of Cell Adhesion with Supervised Training and Learning Environment (CASTLE) workflow. ‘Batch Process’ and ‘Analysis’ are automated in the CASTLE workflow, while ‘Training’ and ‘Validation’ involve manual interaction.

**Table S1:** Table of the validation results on the data produced from the study investigating the role of the protein Gal-9 on leukocyte adhesion. The images automatically processed were compared to a manual analysis by an expert, including the time it took CASTLE and TWS to analyse the images on the computer configuration detailed in Section 2.7. For all iterations, 35 images were validated.

| Validation | Phase | $F_1$ score | Precision | Recall | Time (min:sec) |
| --- | --- | --- | --- | --- | --- |
| Manual | Bright | - | - | - | 26:00 |
|  | Dark | - | - | - | 49:00 |
| Gal-9 study processed by CASTLE<br>(no Transition class) | Bright | 75 | 67 | 84 | 05:08 |
|  | Dark | 87 | 96 | 83 | 05:13 |
| Gal-9 study processed by CASTLE<br>(with Transition class) | Bright | 84 | 80 | 90 | 05:11 |
|  | Dark | 90 | 83 | 98 | 05:17 |
| Gal-9 study processed by TWS<br>(with Transition class) | Bright | 68 | 54 | 93 | 27:00 |
|  | Dark | 58 | 41 | 97 | 40:52 |

**Table S2:** Table of the validation results on the data produced from the study investigating the role of the protein ICAM-1-Fc on leukocyte adhesion. The images automatically processed were compared to a manual analysis by an expert, including the time it took CASTLE to analyse the images on the computer configuration detailed in Section 2.7 For the study investigating the protein ICAM-1, 28 images were analysed and validated. Here, ‘with no additional training’ refers to the same training in CASTLE as that in the study of the role of the protein Gal-9 on leukocyte adhesion while ‘with additional training’ refers to the addition of 1 image from the same study being analysed was added to the training in CASTLE.

| Validation | Phase | $F_1$ score | Precision | Recall | Time (min:sec) |
| --- | --- | --- | --- | --- | --- |
| ICAM-1 study processed by CASTLE<br>(with no additional training) | Bright | 91 | 87 | 96 | 03:59 |
|  | Dark | 73 | 96 | 59 | 04:04 |
| ICAM-1 study processed by CASTLE<br>(with additional training) | Bright | 91 | 89 | 92 | 05:51 |
|  | Dark | 83 | 87 | 80 | 05:29 |

**Table S3:** Table of the validation results on the data produced from the study investigating the role of the protein vWF on platelet adhesion. The images automatically processed were compared to a manual analysis by an expert, including the time it took CASTLE to analyse the images on the computer configuration detailed in Section 2.7. For the study investigating the protein vWF on platelet adhesion, 2 images were validated before completing the analysis on 30 images taken sequentially on the same assay for each condition. Here, ‘with no additional training’ refers to the same training in CASTLE as that in the study of the role of the protein Gal-9 on leukocyte adhesion while ‘with additional training’ refers to training on only 2 images from the study of the role of the protein vWF on platelet adhesion.

| Validation | Phase | $F_1$ score | Precision | Recall | Time (min:sec) |
| --- | --- | --- | --- | --- | --- |
| vWF study processed by CASTLE<br>(with no additional training) | Bright | 68 | 99 | 51 | 08:25 |
|  | Dark | 80 | 77 | 83 | 08:20 |
| vWF study processed by CASTLE<br>(with additional training) | Bright | 89 | 98 | 81 | 08:21 |
|  | Dark | 71 | 59 | 90 | 08:29 |

**Figure S2:**
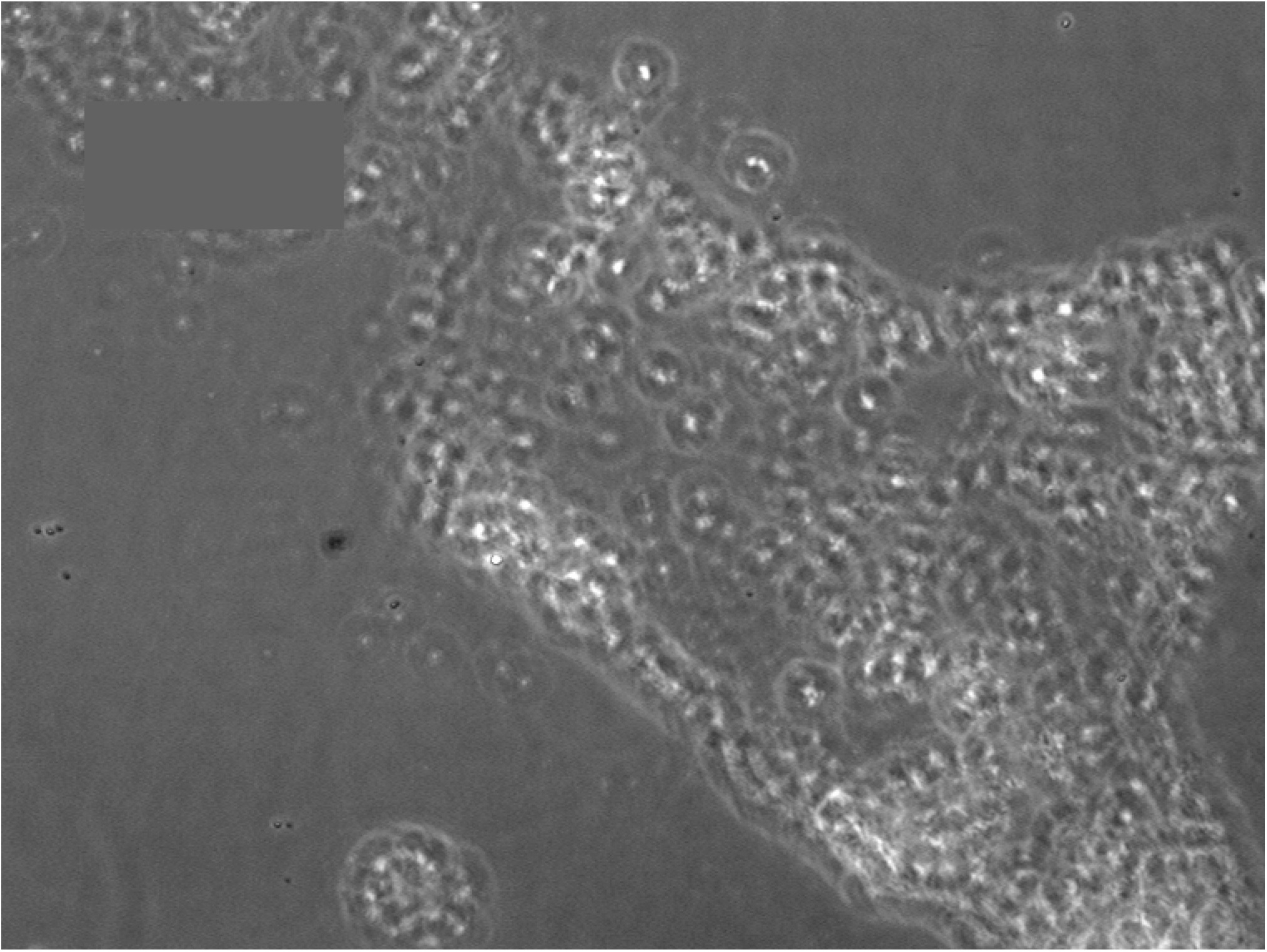
The anomolous image in the PBMC sample group that was exposed to the control BSA protein. We can see that the image gives off the familiar ‘halo’ effect as a neutrophil from irregular particles seemingly trapped under the coating of protein applied to the surface.

**Figure S3:**
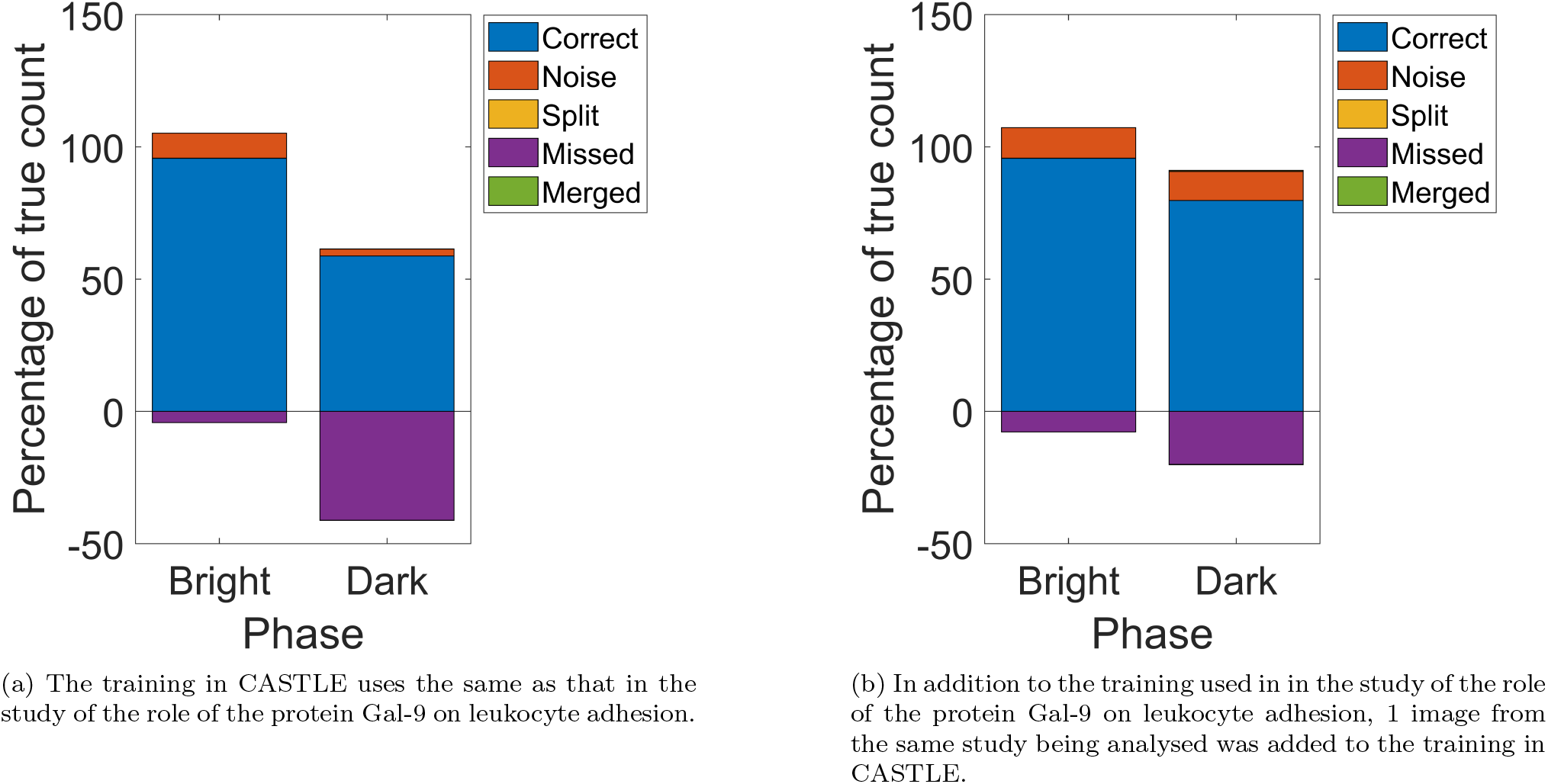
CASTLE validated on the data produced from the study of the role of the protein ICAM-1 on leukocyte adhesion. (a) The training in CASTLE uses the same as that in the study of the role of the protein Gal-9 on leukocyte adhesion. (b) In addition to the training used in in the study of the role of the protein Gal-9 on leukocyte adhesion, 1 image from the same study being analysed was added to the training in CASTLE.

**Figure S4:**
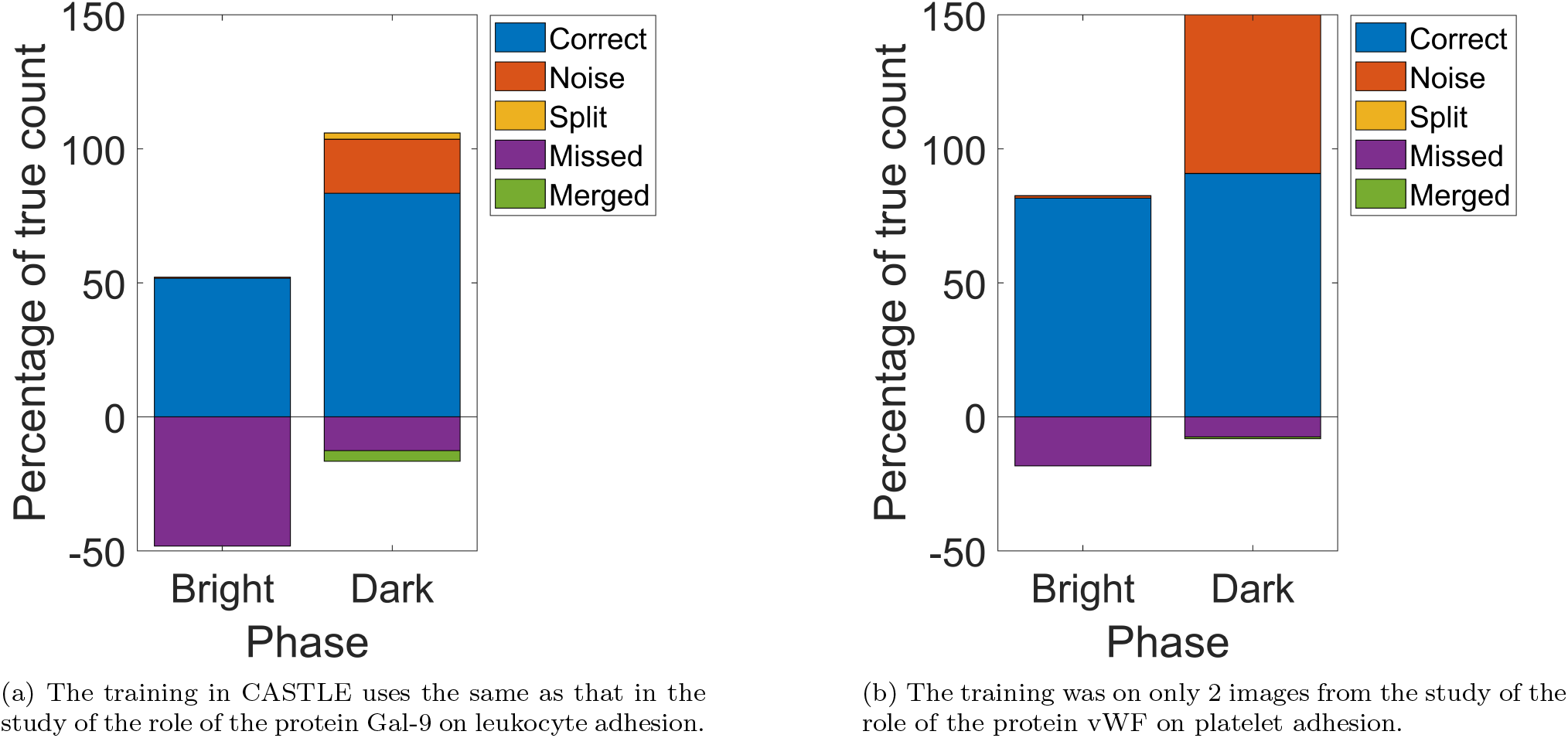
CASTLE validated on the data produced from the study of the role of the protein ICAM-1-Fc on leukocyte adhesion. (a) The training in CASTLE uses the same as that in the study of the role of the protein Gal-9 on leukocyte adhesion. (b) The training was on only 2 images from the study of the role of the protein vWF on platelet adhesion.

**Figure S5:**
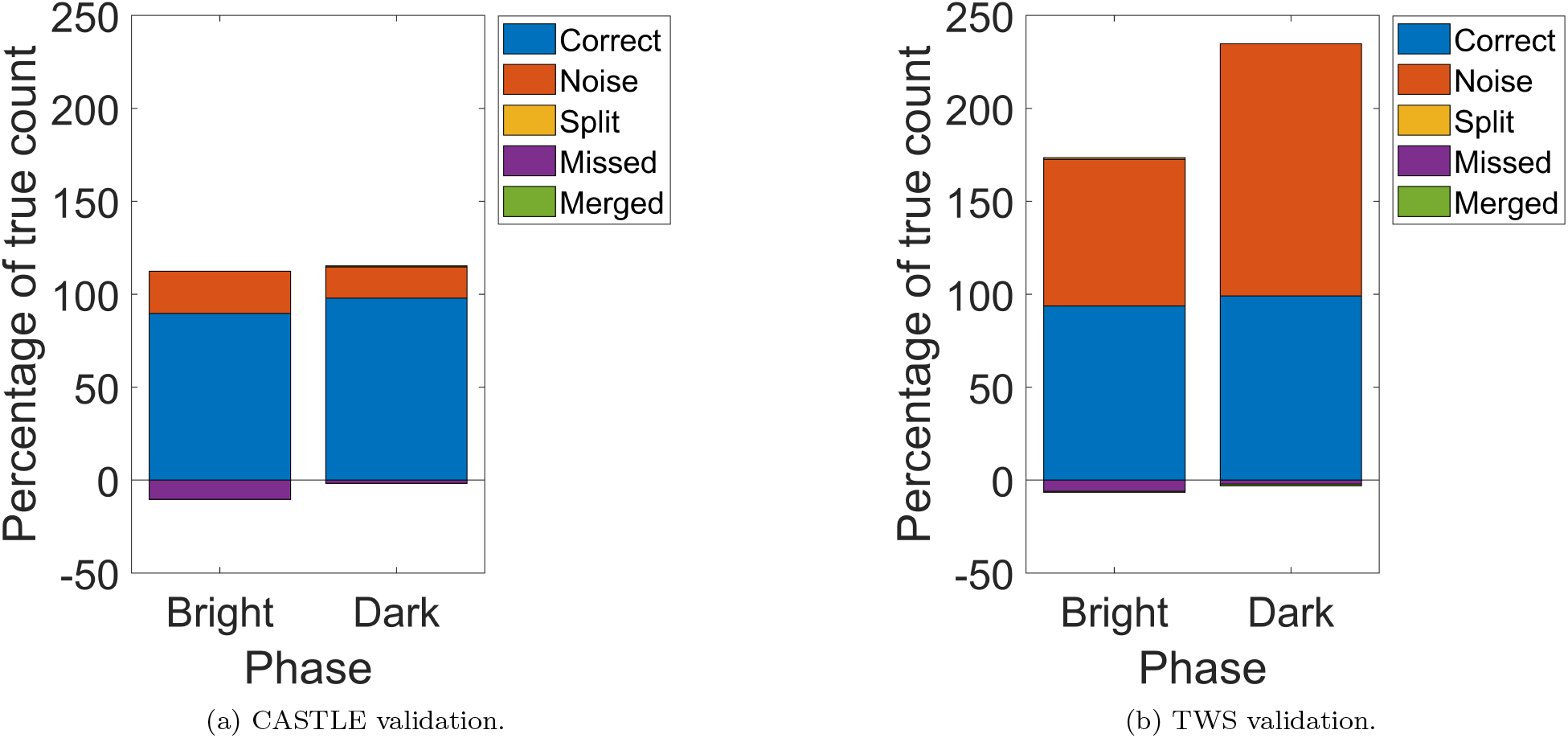
Comparison between CASTLE and TWS on the data produced from the study of the role of the protein Gal-9 on leukocyte adhesion. (a) CASTLE validation. (b) TWS validation. Both were trained with the same set of images with similar inputs for the same number of training classes, including a transition class.

**Table S4:** Hypothesis testing on the difference in means for total cell adhesion per mm^2^ × 10^6^. Here * identifies the comparisons that were statistically significant at the 95% confidence level and ** identifies those that are statistically significant at the 99% confidence level.

| Cell Type | Comparison | p-value |
| --- | --- | --- |
| PBMC | 50 & 100 | 0.285051 |
|  | 50 & BSA | 0.000017 ** |
|  | 100 & BSA | 0.001132 ** |
| PMN | 20 & BSA | 0.000464 ** |

**Table S5:**
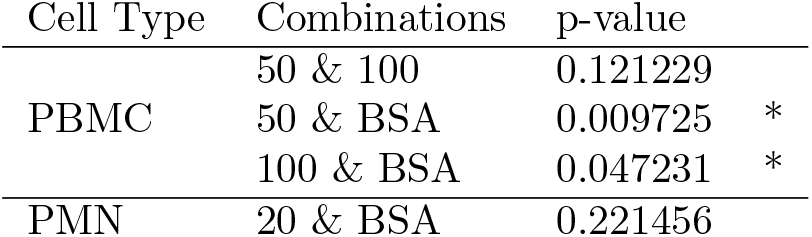
Hypothesis testing on the difference in means for phase bright cell adhesion per mm^2^ × 10^6^. Here * identifies the comparisons that were statistically significant at the 95% confidence level and ** identifies those that are statistically significant at the 99% confidence level.

**Table S6:** Hypothesis testing on the difference in means for phase dark cell adhesion per mm^2^ × 10^6^. Here * identifies the comparisons that were statistically significant at the 95% confidence level and ** identifies those that are statistically significant at the 99% confidence level.

| Cell Type | Combinations | p-value |
| --- | --- | --- |
| PBMC | 50 & 100 | 0.860492 |
|  | 50 & BSA | 0.000321 ** |
|  | 100 & BSA | 0.000181 ** |
| PMN | 20 & BSA | 0.000191 ** |

**Table S7:**
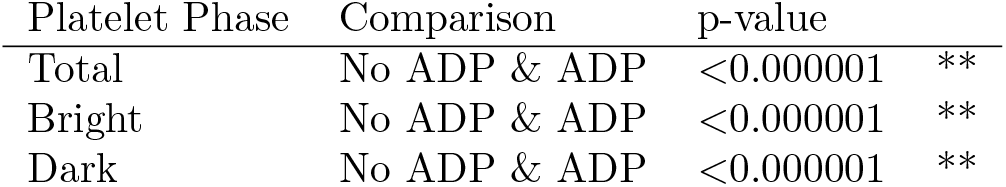
Hypothesis testing on the difference in means for total, phase bright and phase dark adhesion. As before, * identifies the comparisons that were statistically significant at the 95% confidence level and ** identifies those that are statistically significant at the 99% confidence level.

**Figure S6:**
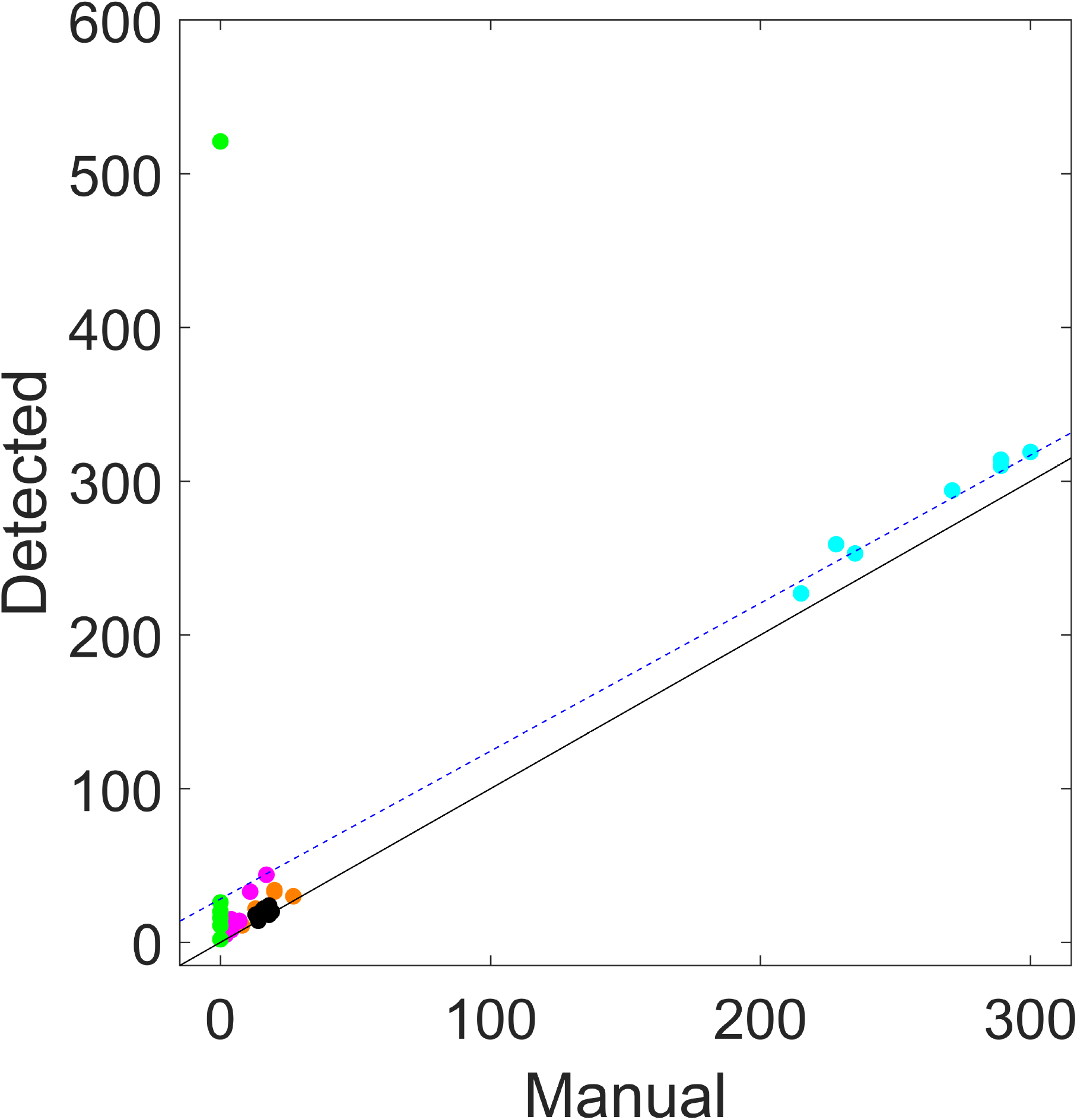
Comparison of automatic to manual detection of adherent phase dark cells for each image in the study into the role of the protein Gal-9 on leukocyte adhesion. Figure 5b, i.e. CASTLE validation with a Transition class during training, but with axes extending the full range to include the anomalous image. The regression line (blue, dashed) shows a prediction for the detected counts across the range of cells in any one image. The line of perfect agreement (black, solid) is the desired outcome with the manual count equal to the automatic count.

**Figure S7:**
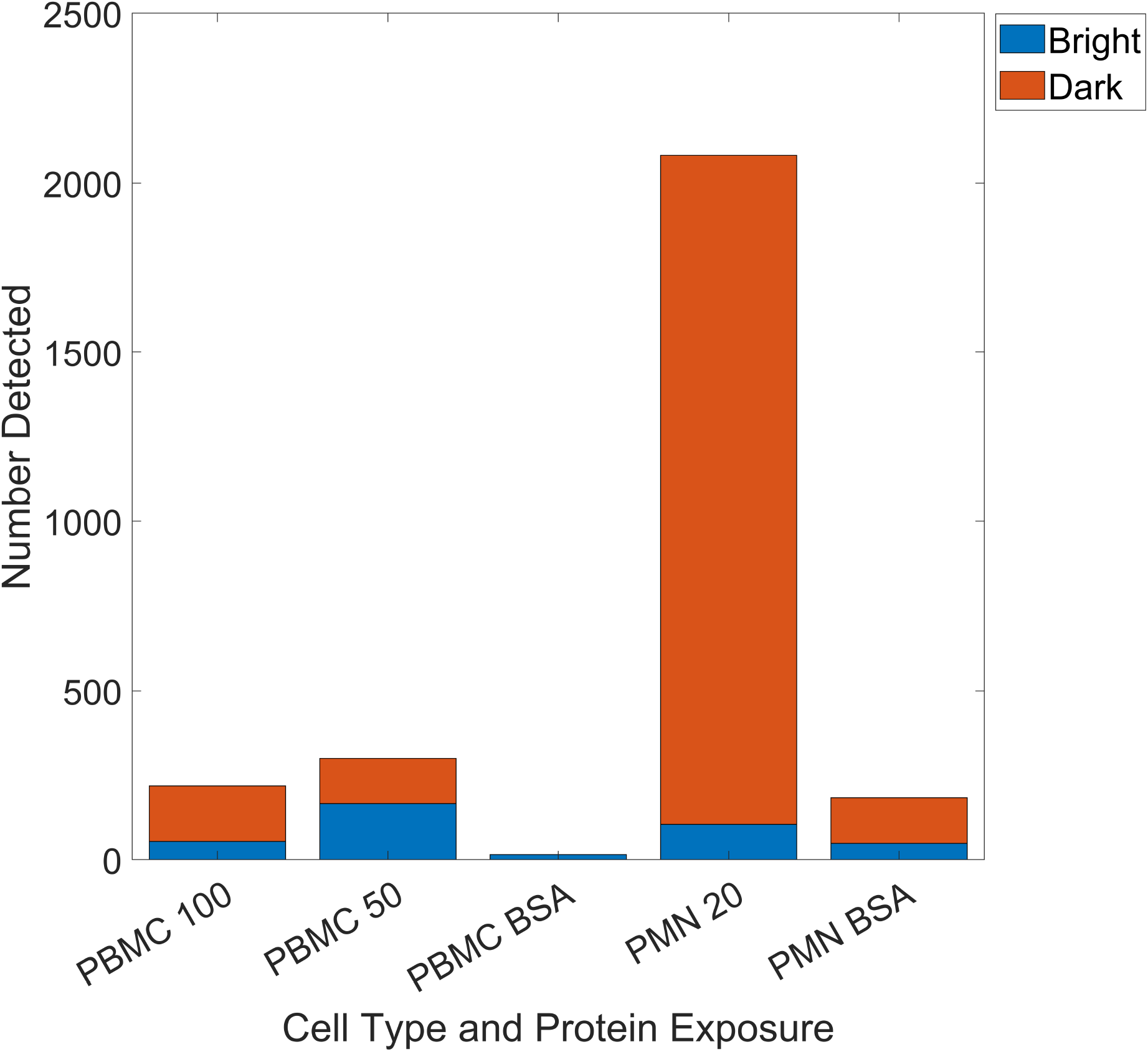
Bar chart displaying the number of leukocytes detected, with the proportion of which are phase bright or dark, by CASTLE for the analysis into the effects of Gal-9, not including the anomalous images.

**Figure S8:**
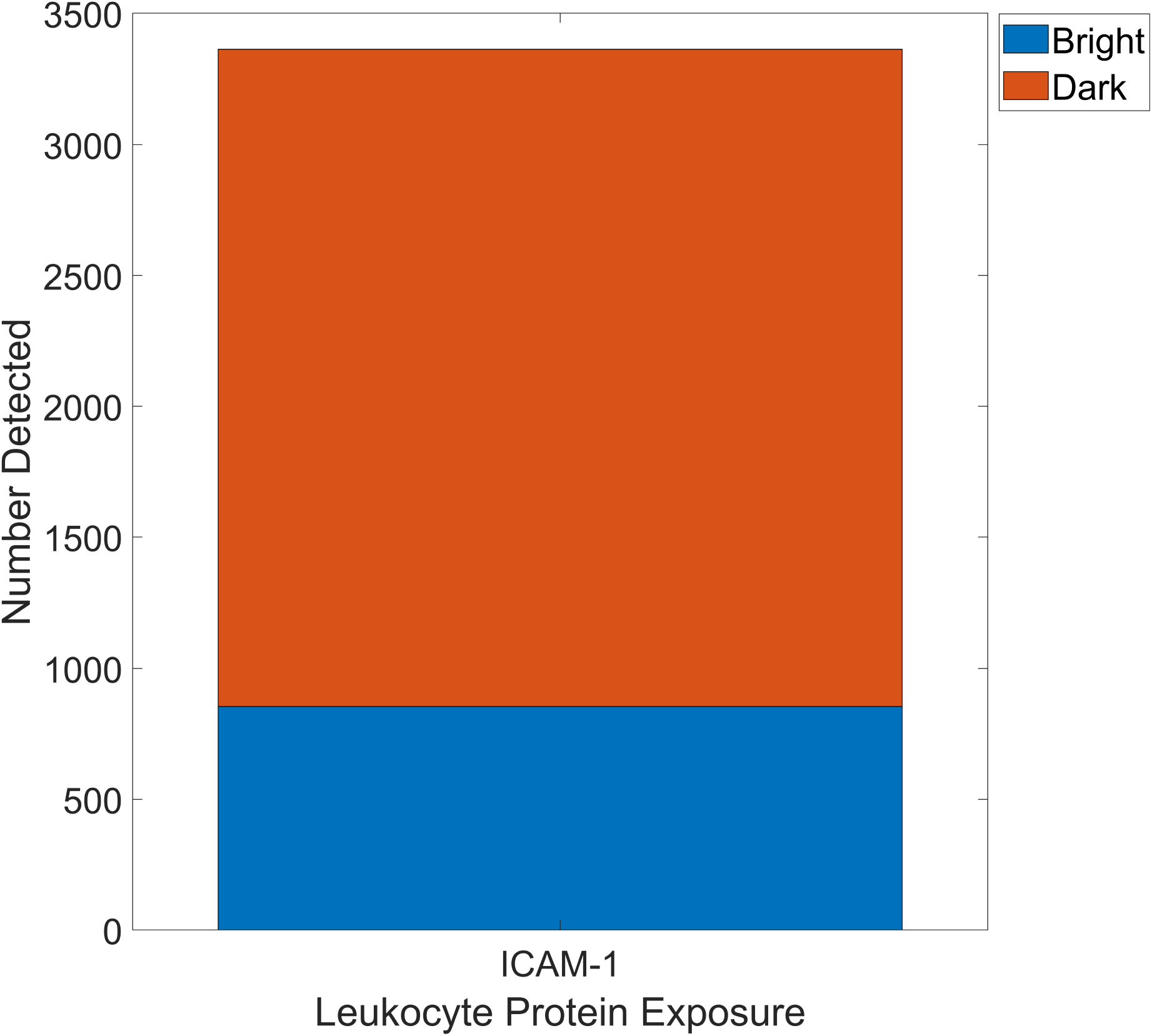
Bar chart displaying the number of leukocytes detected, with the proportion of which are phase bright or dark, by CASTLE for the analysis into the effects of ICAM-1.

**Figure S9:**
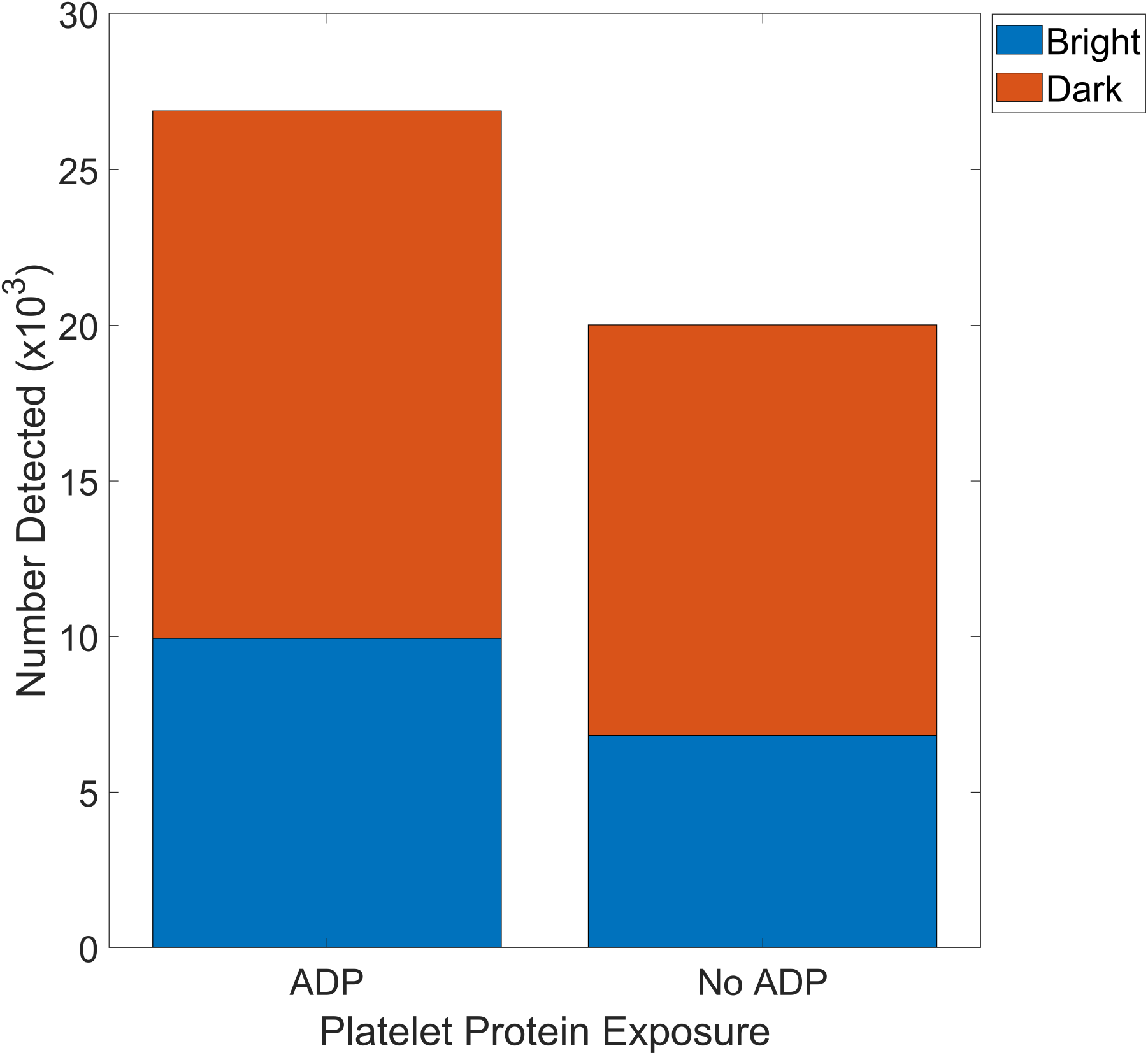
Bar chart displaying the number of platelets detected, with the proportion of which are phase bright or dark, by CASTLE for the analysis into the effects of ADP.

**Figure S10:**
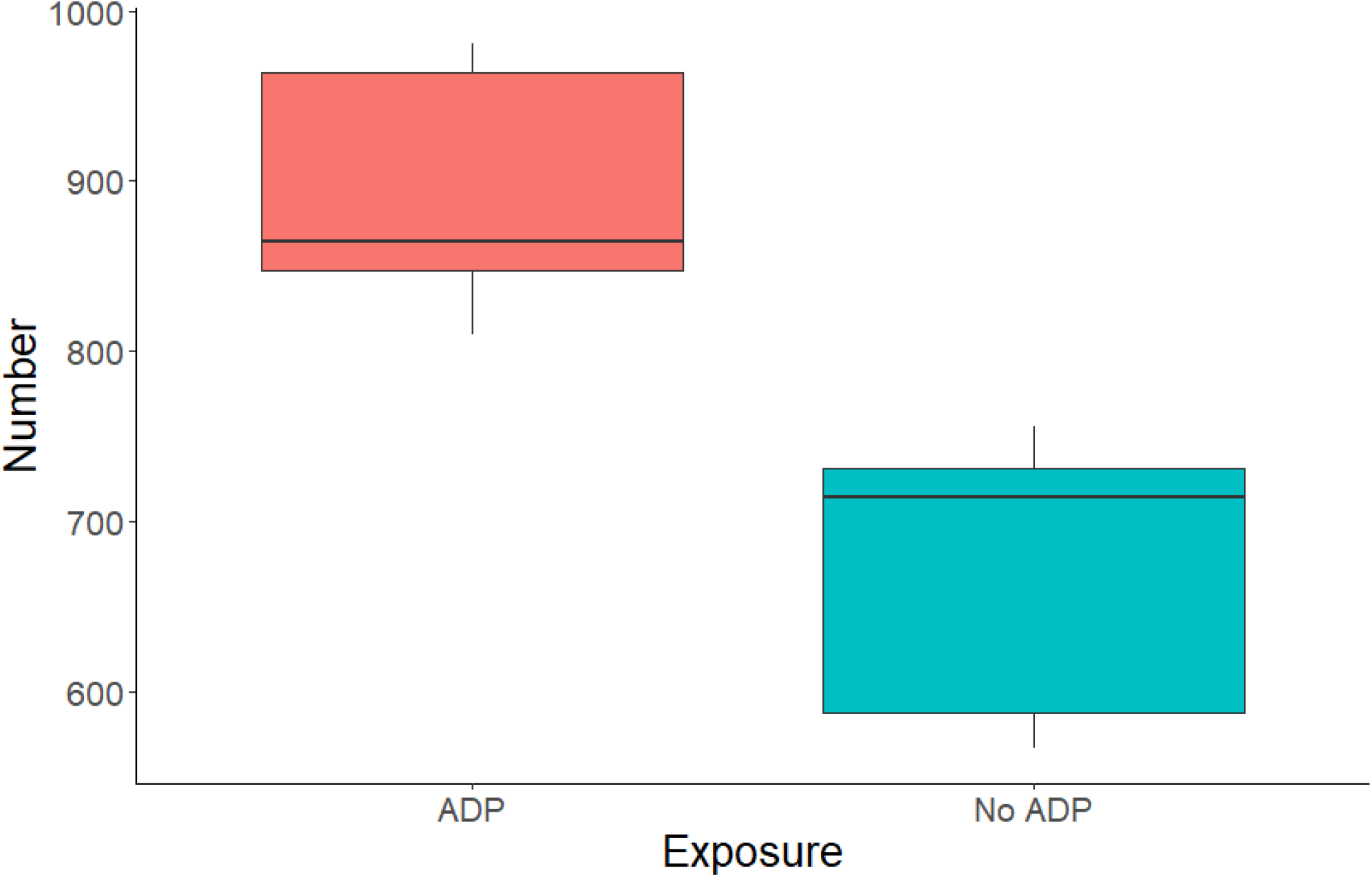
Boxplot for the total number of platelets adhered, in dependence on exposure to ADP.

## Notes

### Competing Interest Statement

The authors have declared no competing interest.

